# GATA1-defective immune-megakaryocytes as possible drivers of idiopathic pulmonary fibrosis

**DOI:** 10.1101/2023.06.20.542249

**Authors:** Francesca Gobbo, Maria Zingariello, Paola Verachi, Mario Falchi, Francesca Arciprete, Fabrizio Martelli, Angelo Peli, Maria Mazzarini, Jeff Vierstra, Carolyn Mead-Harvey, Amylou C. Dueck, Giuseppe Sarli, Stefano Nava, Giacomo Sgalla, Luca Richeldi, Anna Rita Migliaccio

## Abstract

Idiopathic pulmonary fibrosis (IPF) is a progressive fibrotic lung disorder with limited therapeutic options. Insufficient understanding of driver mutations and poor fidelity of currently available animal models has limited the development of effective therapies. Since GATA1 deficient megakaryocytes sustain myelofibrosis, we hypothesized that they may also induce fibrosis in lungs. We discovered that lungs from IPF patients and *Gata1^low^*mice contain numerous GATA1negative immune-poised megakaryocytes that, in mice, have defective RNA-seq profiling and increased TGF-β1, CXCL1 and P-selectin content. With age, *Gata1^low^* mice develop fibrosis in lungs. Development of lung fibrosis in this model is prevented by *P-selectin* deletion and rescued by P-selectin, TGF-β1 or CXCL1 inhibition. Mechanistically, P-selectin inhibition decreases TGF-β1 and CXCL1 content and increases GATA1positive megakaryocytes while TGF-β1 or CXCL1 inhibition decreased CXCL1 only. In conclusion, *Gata1*^low^ mice are a novel genetic-driven model for IPF and provide a link between abnormal immune-megakaryocytes and lung fibrosis.

## Introduction

Idiopathic pulmonary fibrosis (IPF) is a chronic, progressive form of pneumonia associated with fibrosis of unknown etiology^1^. IPF represents the prototypic form of progressive pulmonary fibrosis (PPF), also associated with other non-idiopathic conditions. IPF occurs primarily in older adults (usually >60 years of age) and leads to progressive deterioration of pulmonary function, and eventual death from chronic or acute respiratory failure^1, 2^. Its incidence is 3-9 cases every 100,000 inhabitants and manifests itself prevalently in males with known risk factors such as smoking^3^. Studies of IPF and PPF pathobiology are hampered by limited availability of lung biopsies due to the fact that lung biopsy is no longer needed for diagnosis. While animal models of IPF/PPF have been proposed, the fidelity between the human disease and the currently available models is in general poor. Current animal models employed to study the pathobiology and to identify candidate treatments for IPF are animals (usually mice or rats) forced to breath bleomycin^4^, a chemotherapeutic compound that has lung toxicity as its major adverse effect^5^. Animals subjected to bleomycin develop acute inflammation and fibrosis in the lungs which eventually leads to their death within one-two weeks. In spite of its widespread use, bleomycin has several limitations which reduce its clinical relevance. First, the aggressive progression of fibrosis in this model does not resemble that of IPF, where fibrosis develops slowly over many years. Second, IPF usually affects individuals without previous exposures to chronic inflammatory insults, suggesting that inflammation represents a confounding factor in translating the results from these models to humans. Third and most importantly, the lesions developed in this model are associated with acute alveolar damage, including edema, and have a bronchiolo-centric localization, likely the consequence of the intra-tracheal route of the drug administration^6^. By contrast, in IPF fibrosis occurs mostly in the basal lobes of the lung, is predominantly subpleural, and tends to extend to the para-septal parenchyma generating the typical honeycomb pattern revealed by high-resolution computed tomography (HRCT) of the chest^7^. Such different anatomic localization strongly suggests that the cells responsible for fibrosis in the animal models and in IPF patients are different. Nevertheless, studies in bleomycin mice, validated by observations in patients, have implicated abnormally high levels of several pro-fibrotic proteins, such as transforming growth factor-β (TGF-β)^8^, interleukin-8 (IL-8)^9^, and the adhesion receptor P-selectin (P-SEL)^10^, among others, in the etiology of IPF. In the case of TGF-β, the availability of chemical inhibitors^11^ and validation experiments demonstrating that mice forced to breath adenoviruses containing TGF-β develop fibrosis in the lung^12, 13^, have inspired clinical trials with pirfenidone and nintedanib, two pleiotropic proinflammatory inhibitors also targeting TGF-β, in IPF^14–17^. However, these two drugs have been shown to be effective in slowing down the decline of lung function, but not in preventing disease progression in most IPF patients^14–17^. Therefore, the only cure for IPF is lung transplantation, an option available to only a limited number of patients^1–3^. The clinical needs of IPF patients highlight the need for novel, possibly genetically driven, animal models which more closely resemble the pathological mechanisms of IPF in patients. It is conceivable that these models may also shed light on the genetic drivers for IPF which are still largely unknown. In fact, although familial clustering and association studies suggest that the disease has a genetic basis^3, 18^, a causative genetic link is yet to be established. The identification that loss-of-function mutations in the *telomerase* gene^19^, or in genes, such as *ZCCH8*, which encode proteins required for expression of the *telomerase*^20^, in patients with familiar IPF raised great hope that the genetic cause of the disease had been finally discovered^21^. However, induction of telomerase-deficiency in mice increases the senescence of both hematopoietic and alveolar stem cells but the mice, although more susceptible to bleomycin-induced lung fibrosis, do not develop IPF at baseline^22, 23^. More recently, loss-of-functions mutations in genes encoding surfactant proteins, the expression of which is restricted to alveolar type II cells, have been found in 2-25% of the patients with sporadic and familiar IPF^24^. The pathobiology role of the *SPTFC* mutations in the etiology of IPF was validated by the observation that Tamoxifen-driven conditional knock-in mice harboring one of the *SPTFC* mutations (isoleucine-to-threonine at codon 73 (I73T)) found in the patients develop an aggressive IPF phenotype and die within few weeks from the administration of the drug^25^. However, consideration of possible founder effects suggests that the mutation status (either in telomerase-related or surfactant protein genes), is known for only 4% of all the acquires and congenital cases of IPF^26^. The fact that the driver mutations for >90% of the patients is not known, highlights the need for additional studies to identify other pathobiology pathways that may sustain the disease.

Recent experiments have demonstrated that activation of *c-Jun*, a gene downstream to TGF-β, induces fibrosis in multiple organs, including bone marrow (BM) and lung, in mice^27, 28^. In the case of myelofibrosis (MF), the most severe of the Philadelphia-chromosome negative myeloproliferative neoplasms^29, 30^, c-jun is activated by increased microenvironment bioavailability of TGF-β produced by abnormal megakaryocytes (MK)^31, 32^. In fact, the MK in the BM of patients with myelofibrosis remain immature due to a RSP14 ribosomopathy induced by the driver mutations that reduces the translation of the mRNA for GATA1^33, 34^, the transcription factor essential for their proper maturation^35^. The causative role of this abnormal MK in the etiology of the disease has been validated by experiments demonstrating that mice expressing either the driver mutations or low levels of *Gata1* (*Gata1^low^ mice)* only in MK^36^ develop myelofibrosis with age^37^. The abnormalities of human and murine GATA1 hypomorphic MK include a content greater than normal of TGF-β^38^, CXCL1, the murine equivalent of IL-8^39^, and P-SEL^40^, three of the proinflammatory proteins that have been implicated also in the development of IPF.

MK are important cellular components of the lungs where they exert the function to release platelets (Plt) intravascularly^41, 42^ and to participate in immune reactions against opportunistic infections^43, 44^. In fact, by leveraging single cell RNA-seq and other methods, recent studies have demonstrated the existence of three distinct types of MK, each one exerting a different function^44–46:^ The BM of adult mice and humans contains the Plt-producing, which correspond to the cells previously defined as mature MK, and the niche-supportive and immune MK, both of which have immature morphology. The Plt-producing MK are the most frequent, have high levels of ploidy, express genes related to thrombopoiesis, including the transcription factor GATA1, and are identified by the CD42b/ARNL markers. Niche-supportive MK are mostly >=8N and express high levels of PF4 and TGF-β, and possibly of other pro-inflammatory cytokines, which regulate growth of hematopoietic stem cells, and low levels of GATA1. These cells express the CD61 and MYLK4 markers and may be induced in vitro by thrombopoietin to acquire a mature morphology and to produce Plt. Immune-poised MK are low ploidy with signatures of inflammatory response and myeloid leukocyte activation genes. These cells can be identified by the CD41 and LPS1 markers by confocal microscopy and by the CD41 and CD53 markers by flow cytometry. They have phagocytic activity and can stimulate T cell expansion and have been identified in lungs where their numbers increase in response to infectious challenges and are activated by neutrophil netosis to phagocytize the infective agents^43, 44^. Although previous studies had already implicated MK abnormalities in the development of fibrosis in the lung of bleomycin-treated mice^47^, the pathobiological role of these cells in IPF is far from established. Based on the driving role exerted by GATA1 hypomorphic MK in establishing fibrosis in the BM^33, 34, 37^, we hypothesized that IPF may be driven by a specific MK subpopulation with an abnormally low GATA1 content. Here, we tested this hypothesis by comparing the MK subclasses and their GATA1 content in the lung from patients with IPF and the non-diseased tissue of lung cancer patients and assessed whether lungs from mice carrying the *Gata1*^low^ mutation contain abnormal MK and became fibrotic with age.

## Results

### The lung from patients with IPF contains an excess of GATA1 hypomorphic megakaryocytes biased for the immune lineage

To test the role of MK in the development of IPF, we first compared the frequency and GATA1 content of MK (CD42b^pos^ cells) in lungs from IPF patients with those observed in non-diseased sections from patients with lung cancer (three subject per group) by confocal microscopy (**Figures 1, S1A**). Available clinical information on the patients is described in **Table S1**. Hematoxylin-Eosin and reticulin staining of consecutive sections confirmed abnormal architecture and presence of fibrosis in sections from IPF patients while the overall normal architecture of the sections from No-IPF lungs indicated that they did not contain obvious cancerous tissues (**Figure 1A**). CD42^pos^ MK were identified with *similar* low frequency in sections from IPF and No-IPF lungs. In No-IPF patients, MK were detected inside blood vessels and most of them (80%) contained robust nuclear levels of GATA1 (**Figures 1B, S1A**). By contrast, in IPF-patients, MK were detected within the lung parenchyma in close association with alveolar epithelial cells and <20% of them contained detectable levels of GATA1 (**Figures 1B,C, S1A**). To be noted that, as expected, none of the lung cells which were not stained by CD42b expressed GATA1.

**Figure 1.**
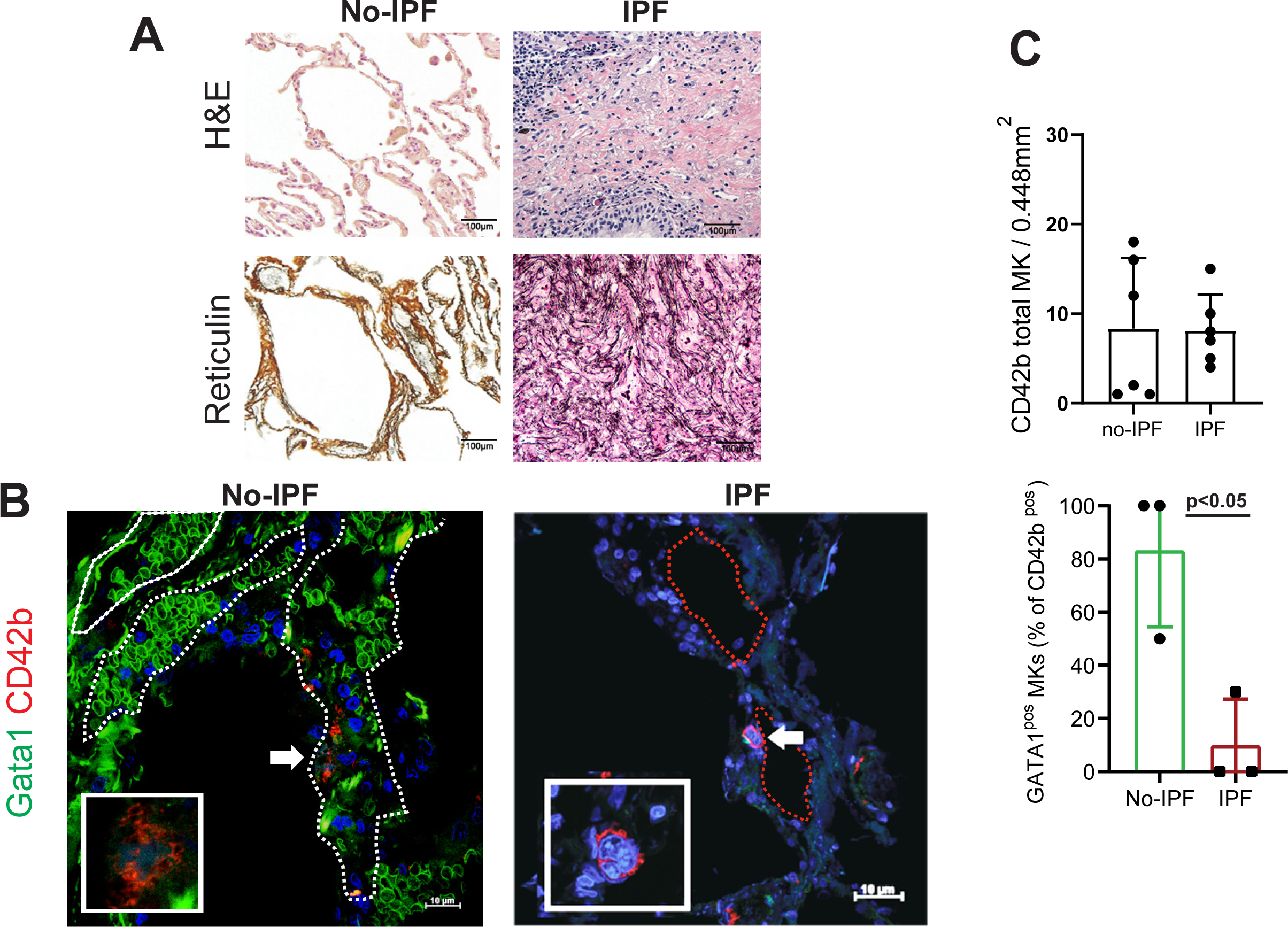
The megakaryocytes in the lung from patients with IPF are hypomorphic for GATA1. A) Hematoxilin&Eosin (H&E) and reticulin staining of the corresponding consecutive sections confirming that the lung from the IPF patient has an overall abnormal architecture with fibrosis. By contrast the archi-tecture of the lung section from the lung cancer patient is overall normal, confirming that we were analyzing a region of the organ free of cancerous cells. Magnification 400x in the panels on the top and 200x in the middle and bottom panels. (B) Double IF staining with anti-GATA1 (green) and CD42b (red, as a marker of MK) of lung sections from one representative IPF patient and one representative no-IPF patient, as indicated. Nuclei are counterstained with DAPI. The insert shows a representative IPF MK at high power fields to show the absence of green signals in its nucleus. The signal captured in the individual channels in additional representative MK are presented in **Figure S1A**. The white and red dotted lines indicate vessels (containing autofluorescent red blood cells) and alveoli (lined by epithelial cells recognized by the morphology of their nuclei, IPF). C) Frequency of CD42b^pos^ MK per area (0.488mm^2^) and percentage of GATA1^pos^ MK (over the total number of CD42b^pos^ cells) observed in the lungs from IPF and no-IPF patients, as indicated. Data are presented as Mean(±SD) and as value in individual subjects (each dot represents a single subject). The data are analyzed by One-way ANOVA and significant p values are indicated within the panels.

The identity of the MK subpopulations present in the lung was assessed by confocal microscopy studies with markers specific for the immune-, HSC-supportive and Plt-poised MK (**Figures 2, S1B**). All MK subpopulations were recognized in the lung from both No-IPF and IPF patients. Again, in No-IPF patients, MK were localized prevalently within blood vessels while in IPF patients they were localized near the alveoli. Interestingly, the frequency of immune-MK was significantly greater (by 2-fold, p<0.05) in lungs from IPF vs. No-IPF patients while that of HSC-supportive and Plt-poised MK was similar in the two groups.

**Figure 2.**
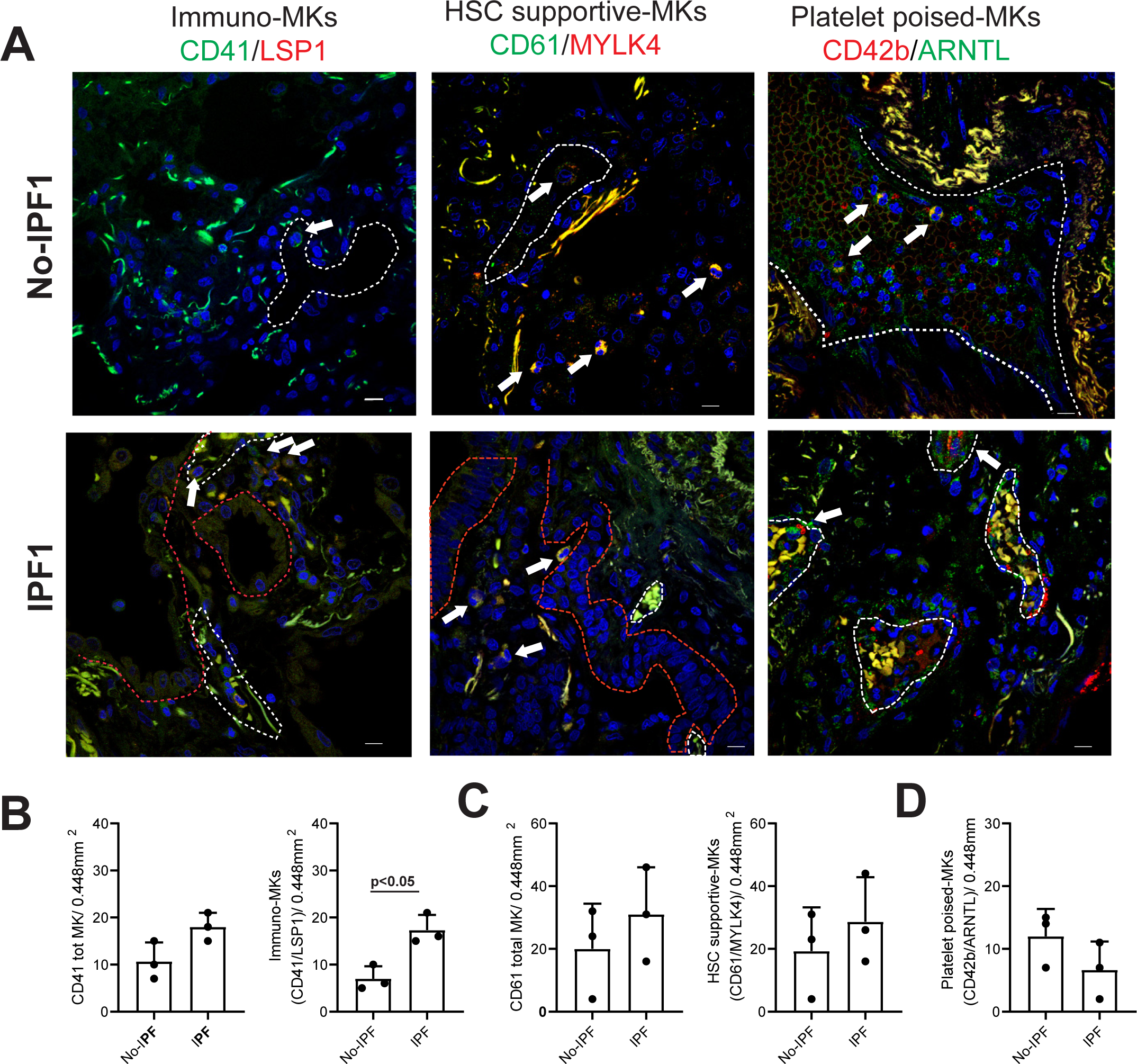
The lung from patients with IPF contains numerous megakaryocytes biased for the immune lineage. A) Confocal microscopy analyses for markers of immune (CD41, green/LSP1, red), HSC-supporting (CD61 green/MYLK4 red) and Plt-poised (CD42b red/ARNTL green) MK in the lung from one representative no-IPF and one IPF patients, as indicated. The signal captured in the individual channels in additional representative MKs are presented in **Figure S1B**. The white and red dotted lines indicate vessels (containing autofluorescent red blood cells) and bronchial epithelium (lined by epithelial cells recognized as cuboidal or columnar cells, IPF). To be noted that the CD41^pos^ cell in the No-IPF section is LSP1^neg^. Original magnification 400x. B-D) Frequency of total CD41^pos^ and CD41^pos^/LPS1^pos^ (immune-MK). D) CD61^pos^ and CD61^pos^/MYLK^pos^ (HSC-supporting MK, E) and total CD42b^pos^ and CD42b^pos^/ARNTL^pos^ (Plt-MK, F) MK detected in the lungs from three No-IPF and three IPF patients, as indicated. Data are presented as Mean(±SD) and as value in individual subjects (each dot represents a single subject). The data are analyzed by One-way ANOVA and significant p values are indicated within the panels.

### Megakaryocytes in the lungs from Gata1^low^ mice have an immature morphology and became biased for the immune-lineage with age

To assess the effects of reduced GATA1 content on the MK subpopulations present in the lung, we compared frequency, maturation state and lineage fate of MK present in mice harboring the *Gata1*^low^ mutation and their wild-type (WT) littermates (**Figures 3,S2,S3** and **Table 1**). MK (CD42b^pos^ cells) are readily identified in lungs from both *Gata1*^low^ and WT littermates by both immune-histochemistry and -fluorescent analysis (**Figures 3A,B, S2A,S3)**. As expected, the nucleus of <10% of CD42b^pos^ cells from the lung of *Gata1^low^* mice contains GATA1 while this transcription factor is clearly detected in the vast majority (>90%) of the MK from the WT littermates (**Figures 3B, S3B**). Also in these mouse studies, expression of GATA1 was restricted to the cells in the lung which express the MK marker CD42b.

**Figure 3.**
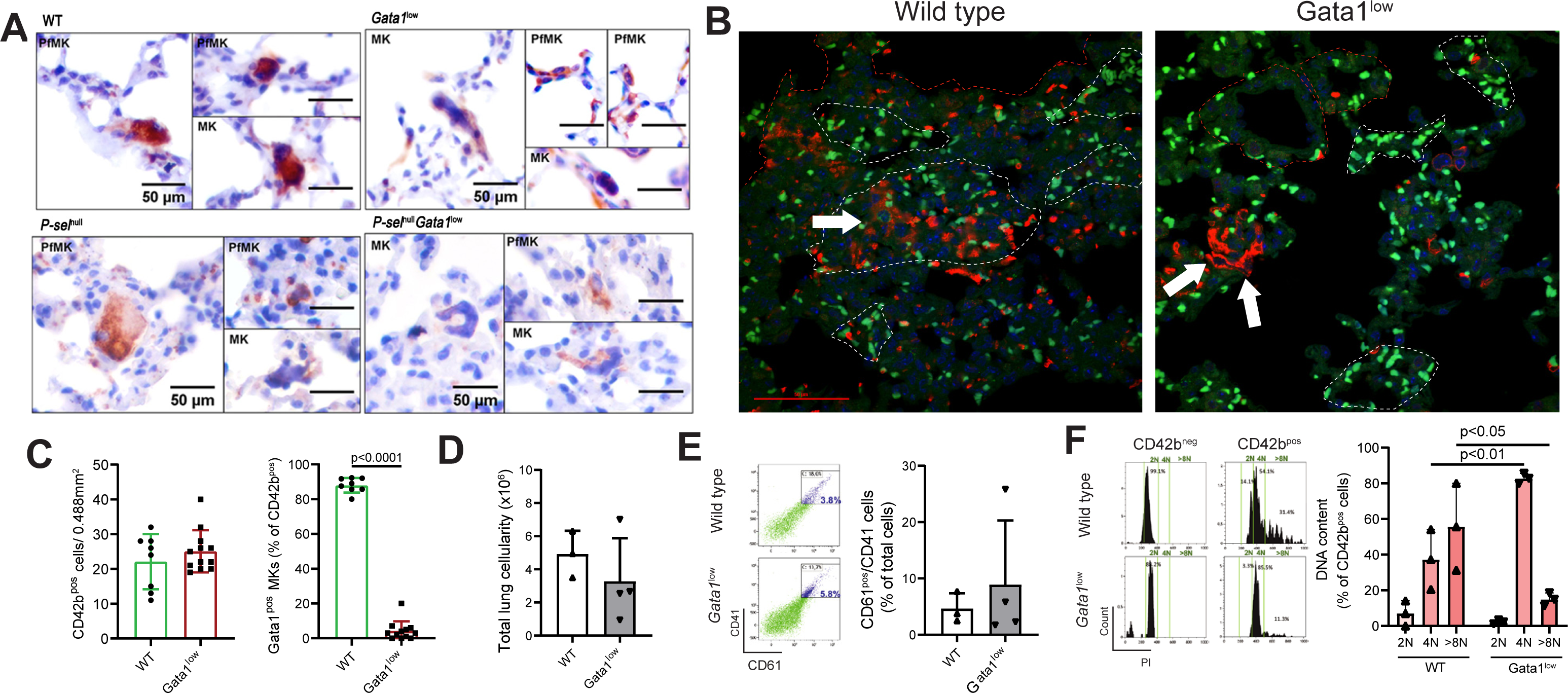
The megakaryocytes in the lungs from Gata1low mice have the morphology of immature cells (reduced platelet territories and low ploidy levels) and are hypomorphic for GATA1. A) IHC with CD42b of lung sections from WT (wild-type) and *Gata1^low^* littermates, as well as from *P-sel^null^* and *P-sel^null^Gata1^low^* mice. The panels present representative MK with immature or mature (presence Plt-forming MK, PfMK) or absence of platelet territories. Scale bar 50µm. Magnification 1000x (See **Figure S2A** for further detail). B) Confocal microscopy with CD42b (red) and GATA1 (green) of MK in the lung from representative WT and *Gata1^low^* littermates. White and red dotted lines indicate vessels (containing autofluorescent red blood cells) and bronchial epithelium (lined by epithelial cells recognized by their canonical morphology). Original Magnification 400x. Frequency of CD42b^pos^ cells/area (0.488mm^2^) and percentage of GATA1^pos^ CD42b^pos^ (over total number of CD42b^pos^ cells) cells observed in lungs from multiple mice are presented. C) Total cell number in lungs of old WT and *Gata1^low^* mice, as indicated. D) Representative flow cytometry analyses with the gate used to determine the frequency of CD41^pos^/CD61^pos^ cells in lungs of WT and *Gata1^low^* mice, as indicated. The frequency observed in multiple experiments is presented on the right. E) Flow cytometry histograms with the level of propidium iodide (PI) incorporated by cells in the CD42b^neg^ (control) and CD42b^pos^ (MK) gate of lung cells from WT and *Gata1^low^* mice. CD42b^neg^ cells were analyzed to determine the staining of cells with a DNA content of 2N. The panel on the right shows the frequency of CD42b^pos^ cells with DNA content 2N, 4N and >8N in lungs from *Gata1^low^* mice and WT littermates (both 15-months old). In B) and E) data are presented as Mean (±SD) and as value in individual mice (each symbol a single mouse) and were analyzed by ANOVA. Statistically significant p values are indicated within the panels.

**Table 1:** Comparison of age, sex, and histological parameters between WT vs *Gata1*^low^ mice and *Gata1*^low^ mice vs P-sel*^null^*Gata1*^low^*.

|  | *Gata1*<sup>low</sup><br>(n=47) | WT<br>(n=18) | Unadjusted<br>p-value | Age-adjusted<br>p-value | Sex-adjusted<br>p-value | P-sel<sup>null</sup>*Gata1*<sup>low</sup><br>(n=25) | Unadjusted<br>p-value | Age-adjusted<br>p-value | Sex-adjusted<br>p-value |
| --- | --- | --- | --- | --- | --- | --- | --- | --- | --- |
| **Age (months)** |  |  |  |  |  |  |  |  |  |
| N (Missing) | 47 (0) | 18 (0) |  |  |  | 25 (0) |  |  |  |
| Mean (±SD) | 12.6 (±5.44) | 8.7 (±5.56) | 0.01<sup>a</sup> |  |  | 10.4 ± (4.04) | 0.07<sup>a</sup> |  |  |
| Median | 12.0 | 10.5 |  |  |  | 11.0 |  |  |  |
| Range | 2.0, 24.0 | 2.0, 18.0 |  |  |  | 3.0, 16.0 |  |  |  |
| **Sex, n (%)** |  |  | 0.37<sup>b</sup> |  |  |  | 0.62<sup>b</sup> |  |  |
| Female | 5 (31.3%) | 5 (31.3.7%) |  |  |  | 14 (56.0%) |  |  |  |
| Male | 11 (68.8%) | 11 (68.8%) |  |  |  | 11 (44.0%) |  |  |  |
| N (Missing) | 45 (2) | 16 (8) |  |  |  | 25 (0) |  |  |  |
| **Total MK in the lung/144mm<sup>2</sup>** |  |  |  |  |  |  |  |  |  |
| N (Missing) | 22 (25) | 9 (9) |  |  |  | 10 (15) |  |  |  |
| Mean (±SD) | 1.5 (±0.91) | 3.6 (±2.42) | 0.001<sup>a</sup> | <0.001<sup>c</sup> | 0.004<sup>c</sup> | 15 (±4.9) | 0.80 | 0.<80<sup>a</sup> | 0.14<sup>c</sup> |
| Median | 1.3 | 3.0 |  |  |  | 1.5 |  |  |  |
| Range | 1.1, 8.3 | 0.0, 3.8 |  |  |  | 8, 22 |  |  |  |
| **Small MK/144mm<sup>2</sup>** |  |  |  |  |  |  |  |  |  |
| N (Missing) | 22 (25) | 9 (9) |  |  |  | 10 (15) |  |  |  |
| Mean (±SD) | 0.7 (±0.84) | 3.4 (±2.53) | <0.001<sup>a</sup> | <0.001<sup>b</sup> | <0.001<sup>c</sup> | 0.8 (±0.39) | 0.70<sup>a</sup> | 0.48<sup>c</sup> | 0.01<sup>c</sup> |
| Median | 0.4 | 3.0 |  |  |  | 0.6 |  |  |  |
| Range | 0.0, 3.6 | 1.0, 8.3 |  |  |  | 0.2, 1.4 |  |  |  |
| **Large MK/144mm<sup>2</sup>** |  |  |  |  |  |  | 0.37<sup>a</sup> | 0.43<sup>c</sup> | 0.72<sup>c</sup> |
| N (Missing) | 22 (25) | 9 (9) |  |  |  | 10 (15) |  |  |  |
| Mean (±SD) | 0.9 (±0.59) | 0.3 (±0.42) | 0.007<sup>a</sup> | 0.009<sup>c</sup> | 0.02<sup>c</sup> | 0.7 (±0.36) | 0.37<sup>a</sup> | 0.43<sup>c</sup> | 0.72<sup>c</sup> |
| Median | 0.7 | 0 |  |  |  | 0.6 |  |  |  |

|  | *Gata1*<sup>low</sup><br>(n=47) | WT<br>(n=18) | Unadjusted<br>p-value | Age-adjusted<br>p-value | Sex-adjusted<br>p-value | P-sel<sup>null</sup>*Gata1*<sup>low</sup><br>(n=25) | Unadjusted<br>p-value | Age-adjusted<br>p-value | Sex-adjusted<br>p-value |
| --- | --- | --- | --- | --- | --- | --- | --- | --- | --- |
| Range | 0.0, 2.3 | 0.0, 1.2 |  |  |  | 0.2, 1.2 |  |  |  |
| **Severity of lung inflammation, n (%)** |  |  | >0.99<sup>b</sup> | 0.85<sup>d</sup> | 0.86<sup>d</sup> |  | 0.27<sup>b</sup> | 0.08<sup>d</sup> | 0.34<sup>d</sup> |
| 0 | 10 (27.0%) | 3 (25.0%) |  |  |  | 9 (50.0%) |  |  |  |
| 1 | 11 (29.7%) | 4 (33.3%) |  |  |  | 4 (22.2%) |  |  |  |
| 2 or 3 | 16 (43.2%) | 5 (41.7%) |  |  |  | 5 (27.8%) |  |  |  |
| N (Missing) | 37 (10) | 12 (6) |  |  |  | 18 (7) |  |  |  |
| **Semiquantitative fibrosis, n (%)** |  |  | 0.01<sup>b</sup> | 0.02<sup>d</sup> | 0.001<sup>d</sup> |  | 0.009<sup>b</sup> | 0.01<sup>d</sup> | <0.001<sup>d</sup> |
| 0 | 8 (27.6%) | 8 (66.7%) |  |  |  | 12 (66.7%) |  |  |  |
| 1 | 9 (31.0%) | 4 (33.3%) |  |  |  | 5 (27.8%) |  |  |  |
| 2 or 3 | 12 (41.4%) | 0 (0.0%) |  |  |  | 1 (5.6%) |  |  |  |
| N (Missing) | 29 (18) | 12 (6) |  |  |  | 18 (7) |  |  |  |
| **Quantitative fibrosis** |  |  |  |  |  |  |  |  |  |
| N (Missing) | 29 (18) | 12 (6) |  |  |  | 18 (7) |  |  |  |
| Mean (±SD) | 12.8 (±11.56) | 3.0 (±3.10) | 0.006<sup>a</sup> | 0.01<sup>c</sup> | 0.002<sup>c</sup> | 5.2 (±5.32) | 0.01<sup>a</sup> | 0.04<sup>c</sup> | 0.002<sup>c</sup> |
| Median | 13.2 | 1.9 |  |  |  | 3.1 |  |  |  |
| Range | 0.0, 45.1 | 0.0, 9.0 |  |  |  | 0.3, 16.3 |  |  |  |

<sup>a</sup>Equal variance two sample t-test p-value; <sup>b</sup>Fisher's exact p-value; <sup>c</sup>Parameter p-value from multiple linear regression with age or sex as covariate; <sup>d</sup>Parameter p-value from ordinal logistic regression with age or sex as covariate.

The localization and maturation state of MK in lung sections from *Gata1^low^* and their WT littermates was determined by morphological and flow cytometry observations (**Figures 3, S2** and **Table 1)**. MK were observed either intravascularly or within the perivascular interstitium in the lungs from both WT and *Gata1*^low^ littermates (**Figure S2A**). Most of the CD42b^pos^ cells in the lungs from WT mice displayed a cytoplasm with clearly divided into Plt territories (Plt-forming MK, PfMKs) and appeared small, probably because they had already released Plt which surrounded them (**Figures 2A,S3A, Table 1**). When detected in the alveolar septa, WT MK were arranged into clusters (**Figure S3A**). By contrast, the MK in the lung from *Gata1*^low^ mice had an immature morphology which included a circular shape and reduced levels of Plt territories (**Figures 3A, S3A, Table 1**). Only a minority of the *Gata1*^low^ MK were PfMKs surrounded by CD42b positive Plt (**Figure S3A).** These *Gata1*^low^ PfMKs were found near blood vessels inside the septa, had a reduced cytoplasm which extended itself between the epithelial cells (**Figure S3A)**.

Statistical analyses of the frequency and maturation state of the MK observed in the lungs from a large cohort of mice indicate that, without controlling for the effects of age or sex, indicated that the difference in the frequency of total, small (both reduced in *Gata1*^low^ mice) and large (increased in *Gata1^low^*mice) MK in the lung from WT and *Gata1*^low^ mice is statistically significant (total p=0.001, small p<0.001, large p=0.007) (**Table 1**).

The morphological observations on the MK maturation state were confirmed by flow cytometry studies on the cell ploidy state determined by CD42b and propidium iodide staining. There was a trend indicating that the lung of *Gata1^low^* mice contain less total cells and more CD61^pos^/CD41^pos^ MK by flow cytometry (**Figure 3C,D**). However, these the interpretation of these data is biased by the extensive manipulation necessary to prepare single cell lung suspensions and by the low number of mice (3-5) analyzed. By flow cytometry, we determined that, as expected^46^, the ploidy of CD42b^pos^ cells in WT lungs ranged from 2N up to 64N with approximately 60% of the cells displaying a DNA content >8N (**Figure 3E**). By contrast, the DNA content of the majority (∼**80**%) of CD42b^pos^ cells from *Gata1^low^*lungs was 4N with few (<20%) MK having a DNA content >8N, confirming that the CD42b cells from the lung of *Gata1^low^* mice are less mature than the WT ones.

In the case of MK, low ploidy cells may represent either Plt-poised MK at the beginning of their maturation process or MK poised to exert immune- or HSC-supporting functions. To assess whether enrichment for low ploidy cells observed in the lung of *Gata1^low^* mice might underly a switch among subpopulations, confocal microscopy studies with markers specific for immune-, HSC-supportive and Plt-poised MK subpopulations were performed (**Figures 4D, S4**). These studies included WT mice as control and analyzed age (young and old) as independent variable. MK with a HSC-supportive or Plt-poised phenotype were recognized in the lungs from both young and old WT and *Gata1^low^* mice at a frequency significantly greater than normal at both ages (**Figures 4,S4**). By contrast, immune-MK were present in robust numbers in the lung from old *Gata1^low^* mice but were barely detectable in those from young *Gata1^low^* mice and from young and old WT mice (**Figures 4,S4**).

**Figure 4.**
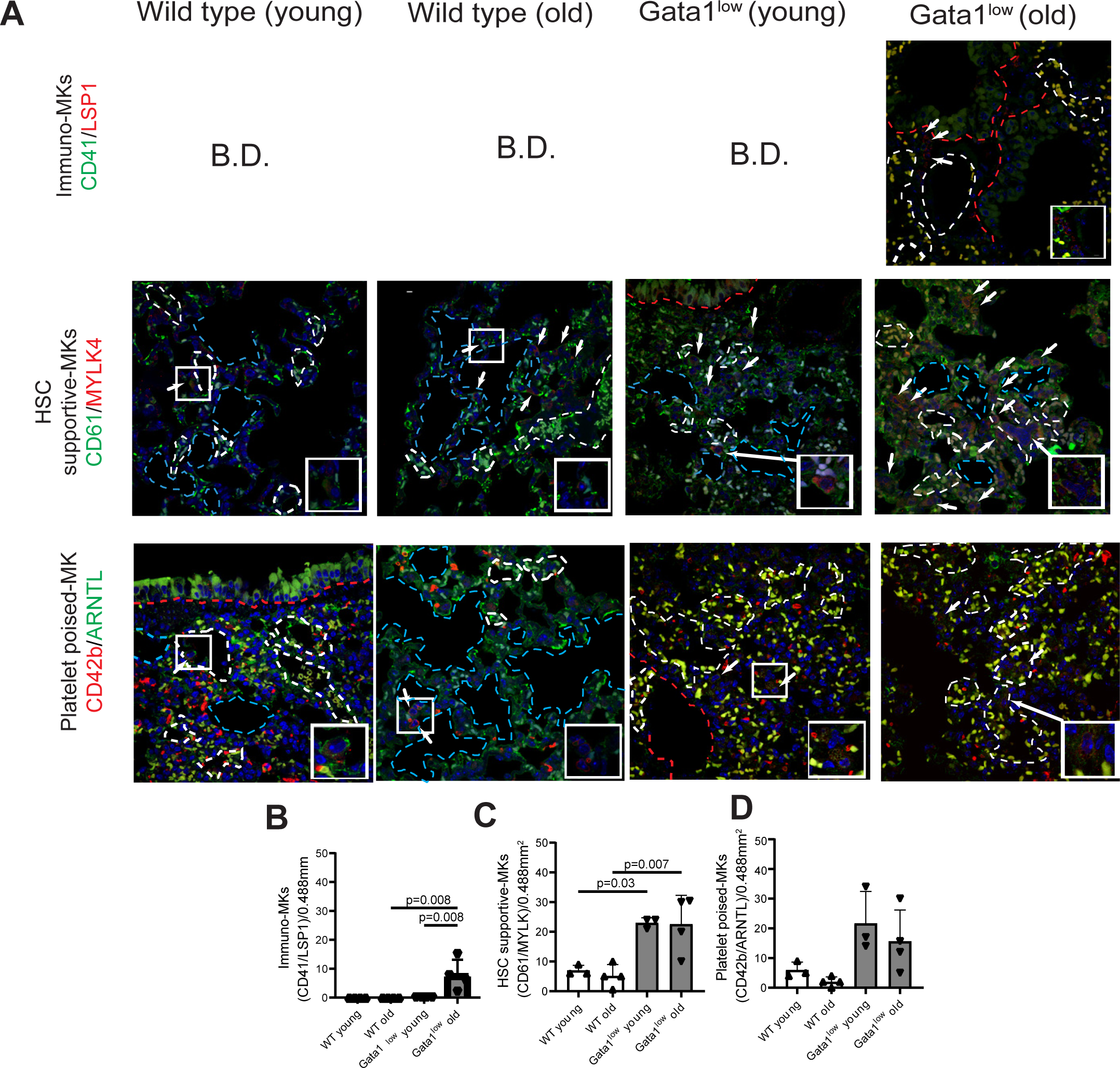
A) Confocal microscopy analyses with markers for the immune (CD41, green/LSP1, red), HSC-supporting (CD61 green/MYLK4 red) and Plt-poised (CD42b red/ARNTL green) MK in the lung from representative WT and *Gata1^low^* mice at 2-3 (young) and 15-months (old) of age, as indicated. MK are indicated by arrows. White dotted lines indicate the contour of blood vessels (recognized by the autofluorescent red blood cells), red dotted lines indicate the profile of the bronchial epithelium and blue dotted lines indicate alveolar spaces contoured by the alveolar epithelium. Original Magnification 400x. The signal from representative MK captured in the individual channels are presented in **Figure S4**. To be noted that Ptl-poised MK are localized within vessels both in the WT and *Gata1^low^* mice. B-D) Frequency of CD41^pos^/LPS1^pos^ (immune-MK), CD61^pos^/MYLK^pos^ (HSC-supporting MK) and CD42b^pos^/ARNTL^pos^ (Plt-poised) MK detected in the lungs from multiple young and old WT and *Gata1^low^*mice. Data are presented as Mean(±SD) and as value in individual subjects (each dot represents a single mouse). The data are analyzed by One-way ANOVA and significant p values are indicated within the panels.

These results indicate that the lungs from old Gata1^low^ mice contain increased levels of immune-MK vs. normal.

### Megakaryocytes from Gata1^low^ mice express a defective immune-signature

To understand the consequences of reduced GATA1 content on the functions of the MK, CD41^pos^ cells (>95% CD41^pos^ by re-analyses) purified from the BM and lung of 15-months old WT and *Gata1*^low^ mice were profiled by RNAseq (**Figures 5, S5**). Principal component analysis (PCA) of the RNAseq data from individual mice indicate a strong separation and clustering among groups, with the differences between tissue of provenience (BM or lung) accounting for approximately 79% of the variance in the expression data (**Figures 5A, S5A**). Overall, the CD41^pos^ cells from the BM and lung of WT mice expressed significantly different levels of 11,828 genes (3,185 upregulated, 8,643 downregulated) while the number of genes significantly expressed at different levels by CD41^pos^ cells from BM and lung of *Gata1^low^*mice were 17,819 (4,848 upregulated, 2,919 downregulated). Comparison of WT and *Gata1^low^* cells from the same origin revealed that the mutation induced extensive changes in gene expression in both organs. The genes significantly different in *Gata1^low^* BM vs WT were 3,257 (1,861 upregulated, 1,396 downregulated) while those in the lung were 2,126 (818 upregulated and 1,308 downregulated) (**Figure S5B**). Interestingly, although both WT and *Gata1^low^*MK were purified according to the CD41 marker, the CD41^pos^ cells from the lung of *Gata1^low^* mice expressed levels of *CD41* (*Itga2b*) significantly greater than those of thee corresponding cells from WT mice (foldchange=2.329, p=0.001), while the expression of *CD61* (*Itgb3*) and *CD42b* (*GpI2b*) was similar between the two groups (**Figure 5B**, **Table 2**), confirming the morphological and flow cytometry observations indicating that the *Gata1^low^* lung from old mice are enriched for immature cells. Comparison of the expression of representative genes characterizing the signatures of the three recently described Mk subpopulations^46^ indicates that most of the expression changes are found among genes that characterize immune-MK (*Clc3*, *Clec4a2*, *Clec4e*, *Clec5a*, *Ctsd*, etc, upregulated and *Cxcl1*, *Ccxl10*, *Il1b*, *Ccl3*, *Cxcl5*, *Lcn2*, etc, downregulated) (**Figure 5B** and **Table 2**). Overall, the abnormalities in the expression signature of MK purified from the BM and lung of *Gata1^low^*mice were similar (**Table 2**), suggesting that they were mostly driven by insufficient expression of the transcription factor rather than by pro-inflammatory cytokines present in the lung microenvironment.

**Figure 5.**
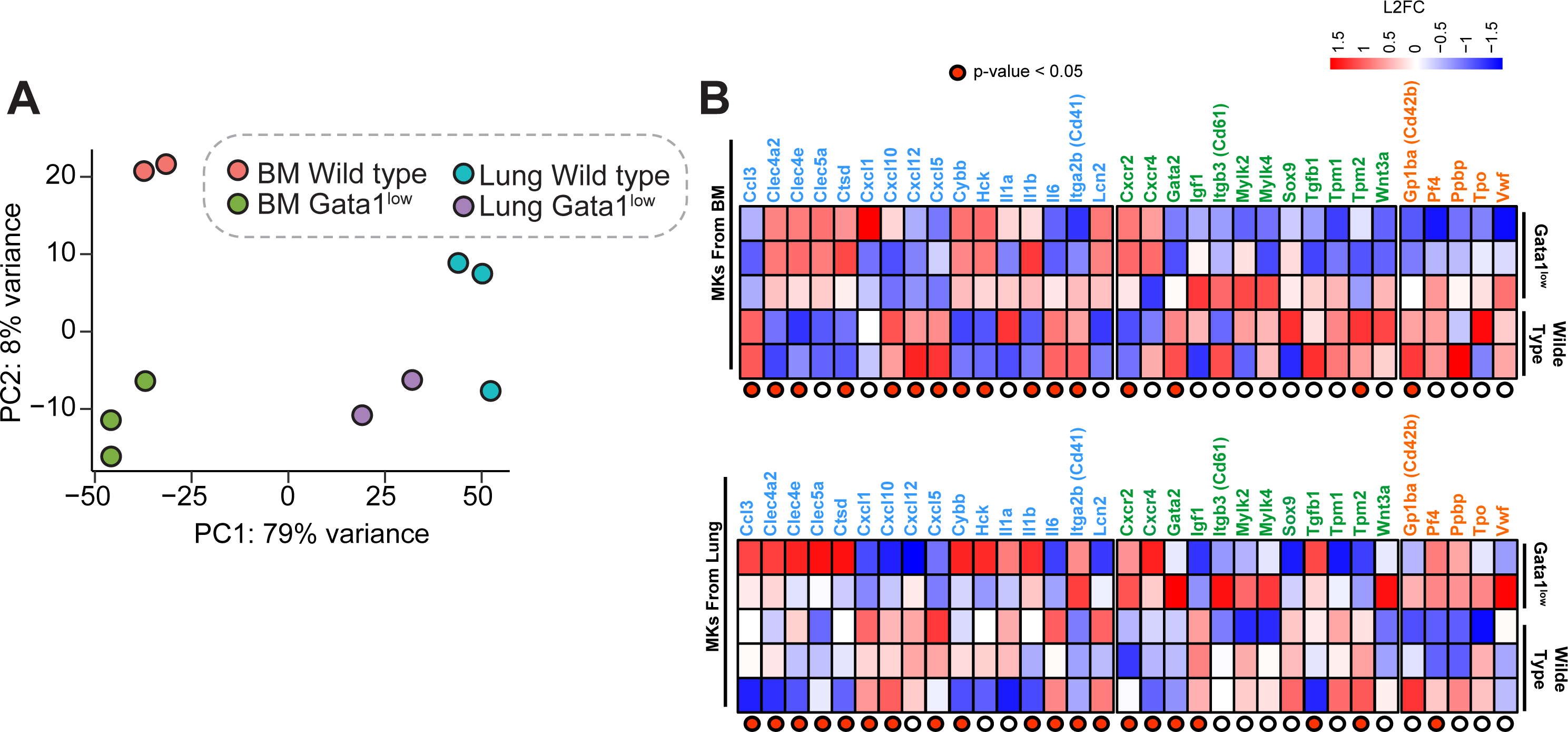
The megakaryocytes from Gata1low mice express a defective immune-signature. A) Principal component analysis (PCA) of the RNAseq data obtained with MK purified from the BM and lungs of individual 15-months old WT and *Gata1^low^* littermates (each dot an individual mouse) as indicated. B) Heatmaps for the level of expression of representative genes which characterize the signature of immune (blue), HSC-supporting (green) and Plt-poised (red) MK described by 43. Statistically significant (p<0.05) differences are indicated by red dots. The significant upregulation of CD41 in the lung of *Gata1^low^*with respect to WT confirms the confocal microscopy data indicating that *Gata1^low^* contain more immune-MK than WT mice. Although immune-MK are higher in number, the overall immune-gene signature of the *Gata1^low^* MK is defective.

**Table 2:**
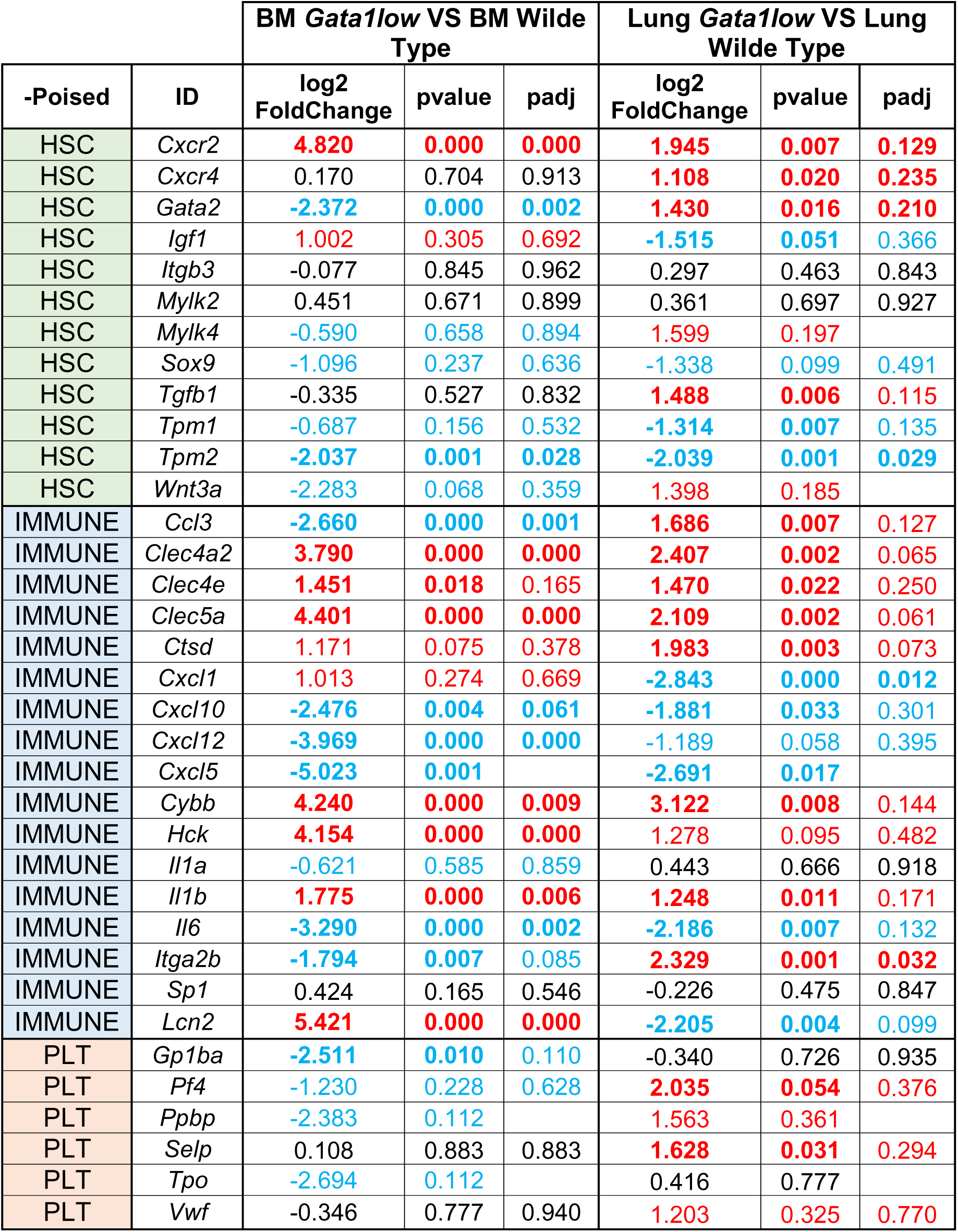
Expression signature for genes which characterize HSC-supportive, immune- and Plt- poised MK populations in CD41pos cells purified from the bone marrow and lung of old *Gata1^low^* mice with respect to the corresponding populations from WT littermates.

| -Poised | ID | BM *Gata1*<sup>low</sup> VS BM Wilde Type |  |  | Lung *Gata1*<sup>low</sup> VS Lung Wilde Type |  |  |
| --- | --- | --- | --- | --- | --- | --- | --- |
|  |  | log2 FoldChange | pvalue | padj | log2 FoldChange | pvalue | padj |
| HSC | *Cxcr2* | **4.820** | **0.000** | **0.000** | **1.945** | **0.007** | **0.129** |
| HSC | *Cxcr4* | 0.170 | 0.704 | 0.913 | **1.108** | **0.020** | **0.235** |
| HSC | *Gata2* | **-2.372** | **0.000** | **0.002** | **1.430** | **0.016** | **0.210** |
| HSC | *Igf1* | **1.002** | **0.305** | **0.692** | **-1.515** | **0.051** | **0.366** |
| HSC | *Itgb3* | -0.077 | 0.845 | 0.962 | 0.297 | 0.463 | 0.843 |
| HSC | *Mylk2* | 0.451 | 0.671 | 0.899 | 0.361 | 0.697 | 0.927 |
| HSC | *Mylk4* | **-0.590** | **0.658** | **0.894** | **1.599** | **0.197** |  |
| HSC | *Sox9* | **-1.096** | **0.237** | **0.636** | **-1.338** | **0.099** | **0.491** |
| HSC | *Tgfb1* | -0.335 | 0.527 | 0.832 | **1.488** | **0.006** | **0.115** |
| HSC | *Tpm1* | **-0.687** | **0.156** | **0.532** | **-1.314** | **0.007** | **0.135** |
| HSC | *Tpm2* | **-2.037** | **0.001** | **0.028** | **-2.039** | **0.001** | **0.029** |
| HSC | *Wnt3a* | **-2.283** | **0.068** | **0.359** | **1.398** | **0.185** |  |
| IMMUNE | *Ccl3* | **-2.660** | **0.000** | **0.001** | **1.686** | **0.007** | **0.127** |
| IMMUNE | *Clec4a2* | **3.790** | **0.000** | **0.000** | **2.407** | **0.002** | **0.065** |
| IMMUNE | *Clec4e* | **1.451** | **0.018** | **0.165** | **1.470** | **0.022** | **0.250** |
| IMMUNE | *Clec5a* | **4.401** | **0.000** | **0.000** | **2.109** | **0.002** | **0.061** |
| IMMUNE | *Ctsd* | **1.171** | **0.075** | **0.378** | **1.983** | **0.003** | **0.073** |
| IMMUNE | *Cxcl1* | **1.013** | **0.274** | **0.669** | **-2.843** | **0.000** | **0.012** |
| IMMUNE | *Cxcl10* | **-2.476** | **0.004** | **0.061** | **-1.881** | **0.033** | **0.301** |
| IMMUNE | *Cxcl12* | **-3.969** | **0.000** | **0.000** | **-1.189** | **0.058** | **0.395** |
| IMMUNE | *Cxcl5* | **-5.023** | **0.001** |  | **-2.691** | **0.017** |  |
| IMMUNE | *Cybb* | **4.240** | **0.000** | **0.009** | **3.122** | **0.008** | **0.144** |
| IMMUNE | *Hck* | **4.154** | **0.000** | **0.000** | **1.278** | **0.095** | **0.482** |
| IMMUNE | *Il1a* | **-0.621** | **0.585** | **0.859** | **0.443** | **0.666** | **0.918** |
| IMMUNE | *Il1b* | **1.775** | **0.000** | **0.006** | **1.248** | **0.011** | **0.171** |
| IMMUNE | *Il6* | **-3.290** | **0.000** | **0.002** | **-2.186** | **0.007** | **0.132** |
| IMMUNE | *Itga2b* | **-1.794** | **0.007** | **0.085** | **2.329** | **0.001** | **0.032** |
| IMMUNE | *Sp1* | 0.424 | 0.165 | 0.546 | -0.226 | 0.475 | 0.847 |
| IMMUNE | *Lcn2* | **5.421** | **0.000** | **0.000** | **-2.205** | **0.004** | **0.099** |
| PLT | *Gp1ba* | **-2.511** | **0.010** | **0.110** | -0.340 | 0.726 | 0.935 |
| PLT | *Pf4* | **-1.230** | **0.228** | **0.628** | **2.035** | **0.054** | **0.376** |
| PLT | *Ppbp* | **-2.383** | **0.112** |  | **1.563** | **0.361** |  |
| PLT | *Selp* | 0.108 | 0.883 | 0.883 | **1.628** | **0.031** | **0.294** |
| PLT | *Tpo* | **-2.694** | **0.112** |  | 0.416 | 0.777 |  |
| PLT | *Vwf* | -0.346 | 0.777 | 0.940 | **1.203** | **0.325** | **0.770** |

Collectively, these results indicate that the immune-MK from the BM and lung of old *Gata1^low^* mice are functionally corrupted.

### Both WT and Gata1^low^ mice are prone to display inflammation in lung with age

Since comparative studies on the baseline pathologies expressed by mice of different strains indicate that CD1 mice, the background where the *Gata1^low^* mutation is hosted, suffer from acute inflammatory pathology, including pneumonia^48^, we first determined whether the presence of the *Gata1^low^* mutation would exacerbate the baseline inflammation that is expected to be present in the lung from CD1 mice as they age (**Figures 6,S2** and **Table 1**).

**Figure 6.**
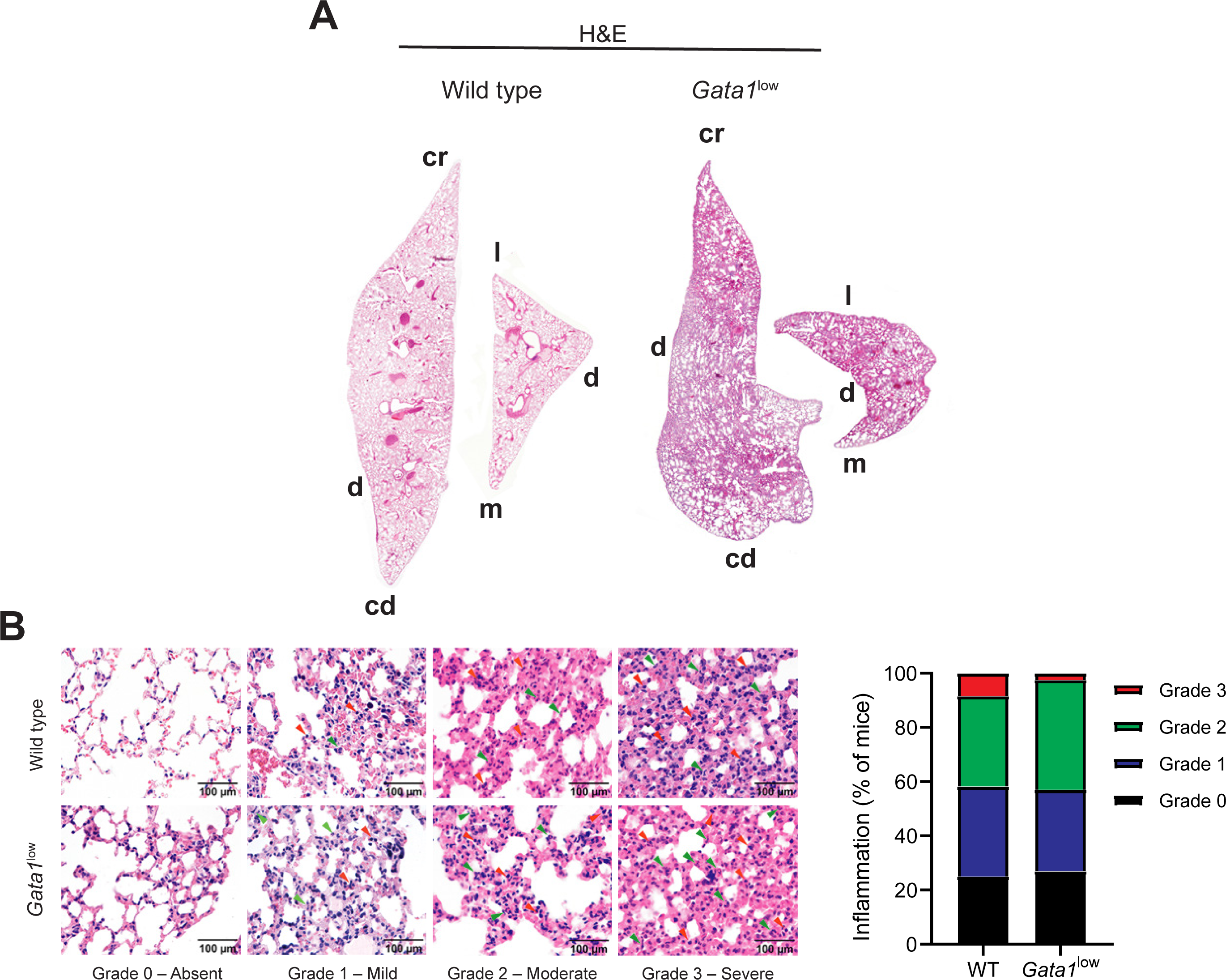
Inflammation is present in lungs from wild-type and Gata1low littermates alike. (A) Stack images of the lungs from representative WT and *Gata1^low^* littermates (both 20 months old) stained by H&E or Masson’s trichrome, as indicated. Original magnification 40x. Abbreviations: cr=cranial: cd=caudal; d=dorsal; l=lateral and m=medial. B) H&E staining of sections of the lungs from representative WT and *Gata1^low^* littermates expressing interstitial pneumonia at grade 0, 1, 2 and 3, as indicated. Red and green arrows indicate lymphocytes and neutrophils respectively. Magnification 200x. The frequency of mice expressing the different grades of inflammation in the two groups is indicated on the right. (see **Table 1** for further detail).

Histological features consistent with interstitial pneumonia are observed in the lungs from a significant number (>40%) of both the WT and *Gata1*^low^ mice analyzed (**Figures 6, S2B,** and **Table 1**). More specifically, interstitial pneumonia (mild, moderate and severe) characterized by infiltration of lymphocytes, macrophages and neutrophils in the alveolar septae and in the peri-bronchial-vascular spaces, was observed in 9 out of 12 WT mice analyzed and in 27 out of 37 of the *Gata1*^low^ mice (**Figure 6**, **Table 1**). In *Gata1*^low^ mice, pneumonia extended from the alveolar septae to the subpleural interstitium (**Figure S2B**). Pneumonia was associated with the presence of infiltrating neutrophils that were often found surrounding the MK (**Figure S2C**). Without controlling for the effects of age or sex, there are no statistically significant differences in severity of lung inflammation between the two genotypes (**Table 1**). Sensitivity analyses adjusting for age and for sex also failed to find a statistically significant difference (**Table 1**).

These data indicate that the presence of the *Gata1^low^* mutation does not exacerbate the baseline interstitial pneumonia developed by mice of the CD1 strain.

### Fibrosis is present only in the lung from old Gata1^low^ mice

Although the fibrosis in the lung from bleomycin-treated animal models is associated with an inflammatory state induced by the chemical insult, it is debated whether inflammation has a direct causative role in the development of IPF. Since the lung from *Gata1^low^*and WT littermates display similar levels of inflammation, we hypothesized that a comparison between the level of fibrosis developed in the lungs of the two strains might clarify whether in IPF inflammation and fibrosis are associated.

In contrast to the lack of difference in inflammation between the two strains, there was an elevated incidence of fibrosis detected by both semiquantitative and quantitative criteria in the lungs from *Gata1*^low^ mice while fibrosis was seldomly observed in those from their WT littermates (**Figures 7,S6,S7A** and **Table 1**). In fact, both by reticulin and Masson’s trichrome staining, fibrosis is not detectable (66.7%) or is overall mild (Grade 1, 33.3%) in the lungs from WT mice. By contrast, it is absent only in a minority (27.6%) of the lungs from *Gata1*^low^ mice and is present at mild (Grade 1) and moderate/severe (Grade 2-3) levels in 41.4% of the mice (**Table 1**) with a penetrance of 85.7% at >22 months of age. Of note, both by Masson staining and immunofluorescent-microscopy with Col 1A and Col III antibodies, MK were consistently found embedded within areas of collagen deposition (**Figure S7**).

**Figure 7.**
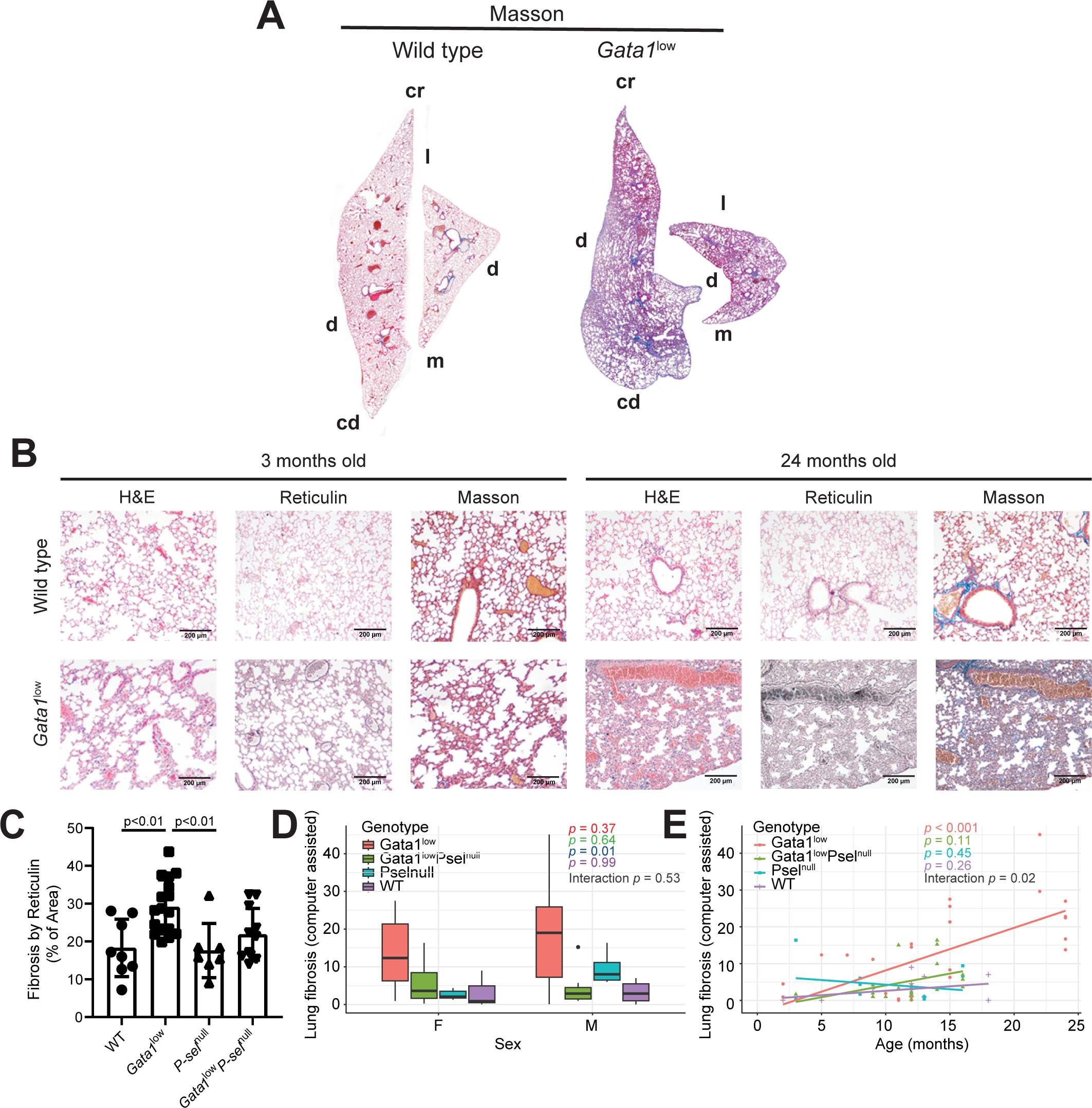
Fibrosis is present only in lungs from old *Gata1^low^* mice. (A) Stack images of the lungs from representative WT and *Gata1^low^* littermates (both 20 months old) stained by Masson’s trichrome, as indicated (consecutive sections of the images presented in Figure 6). Original magnification 40x. Abbreviations: cr=cranial: cd=caudal; d=dorsal; l=lateral and m=medial. B) H&E, Reticulin and Masson’s trichrome staining of consecutive sections from the lungs of representative WT and *Gata1^low^* littermates at 3- and 24-months of age, as indicated. Magnification 100x. C-E) Quantification of fibrosis by reticulin staining. (C) and association of fibro *Gata1^low^*sis in the lung evaluated by Masson’s trichrome staining with sex (D) and age (E), stratified by WT, *Gata1^low^*, *P-sel^null^* and *Gata1^low^P-sel^null^* littermates, as indicated. P-values from One-way ANOVA in C and logistic (D) and simple linear (E) regression models, stratified by genotype are overlayed, along with p-value for the interaction term from a multivariable model including sex and genotype (D) or age and genotype (E) as covariates (see **Table 1** for further detail).

Statistical analyses with a large cohort of mice indicate that, without controlling for the effects of age or sex, there are statistically significant differences between WT and *Gata1*^low^ mice in the level of lung fibrosis as assessed by semiquantitative (*p*=0.01) or by quantitative methods (*p*=0.006) (**Table 1**). In multivariable analysis, the formal test of interaction between genotype and age was statistically significant (*p*=0.02), indicating that age is a statistically significant modifier on the relationship between genotype and lung fibrosis. By contrast, there is insufficient evidence to conclude there is a relationship between sex and fibrosis grade (*p*=0.99) (**Figure 7D,E**).

These data indicate that fibrosis in the lungs is an age-associated trait in mice harboring the *Gata1*^low^ genetic lesion.

### Lungs from old WT and Gata1^low^ mice contain elevated levels of P-SEL, TGF-***β***1 and CXCL1 which only in Gata1^low^ mice are highly expressed by megakaryocytes

Previous results in bleomycin mice have indicated that IPF is associated with greater than normal expression of several pro-inflammatory factors including P-SEL (high in endothelial cells); TGF-β (produced by multiple cell types including alveolar macrophages, epithelial and endothelial cells, interstitial fibroblasts and alveolar Type II cells); and CXCL1, the murine equivalent of human interleukin-8 (IL-8) in bronchoalveolar lavages^10, 49–59^. High levels of TGF-β are also expressed by the alveolar Type II cells of the Tamoxifen-driven conditional knock-in *SPTFCI73T* mouse model^25^. However, the underlying inflammatory state present in the lung of these models hampers assessing which cell type is responsible to produce the pro-inflammatory cytokines responsible for fibrosis. We hypothesized that this assessment may be facilitated by investigations in our model that discriminates between inflammatory (present in both WT and *Gata1^low^* lungs) and fibrotic (*Gata1^low^*lungs only) events. To this aim, we compared the content of P-SEL, TGF-β1 and CXCL1 in the lungs from *Gata1*^low^ and WT littermates by immunohistochemistry (**Figure 8**). Strikingly, the lungs from WT mice contained robust levels of both TGF-β1 and CXCL1 while P-SEL was barely detectable. The levels of both cytokines were 20-fold and 2-fold greater, respectively, and P-SEL was expressed at robust levels in the lungs from *Gata1*^low^ mice.

**Figure 8.**
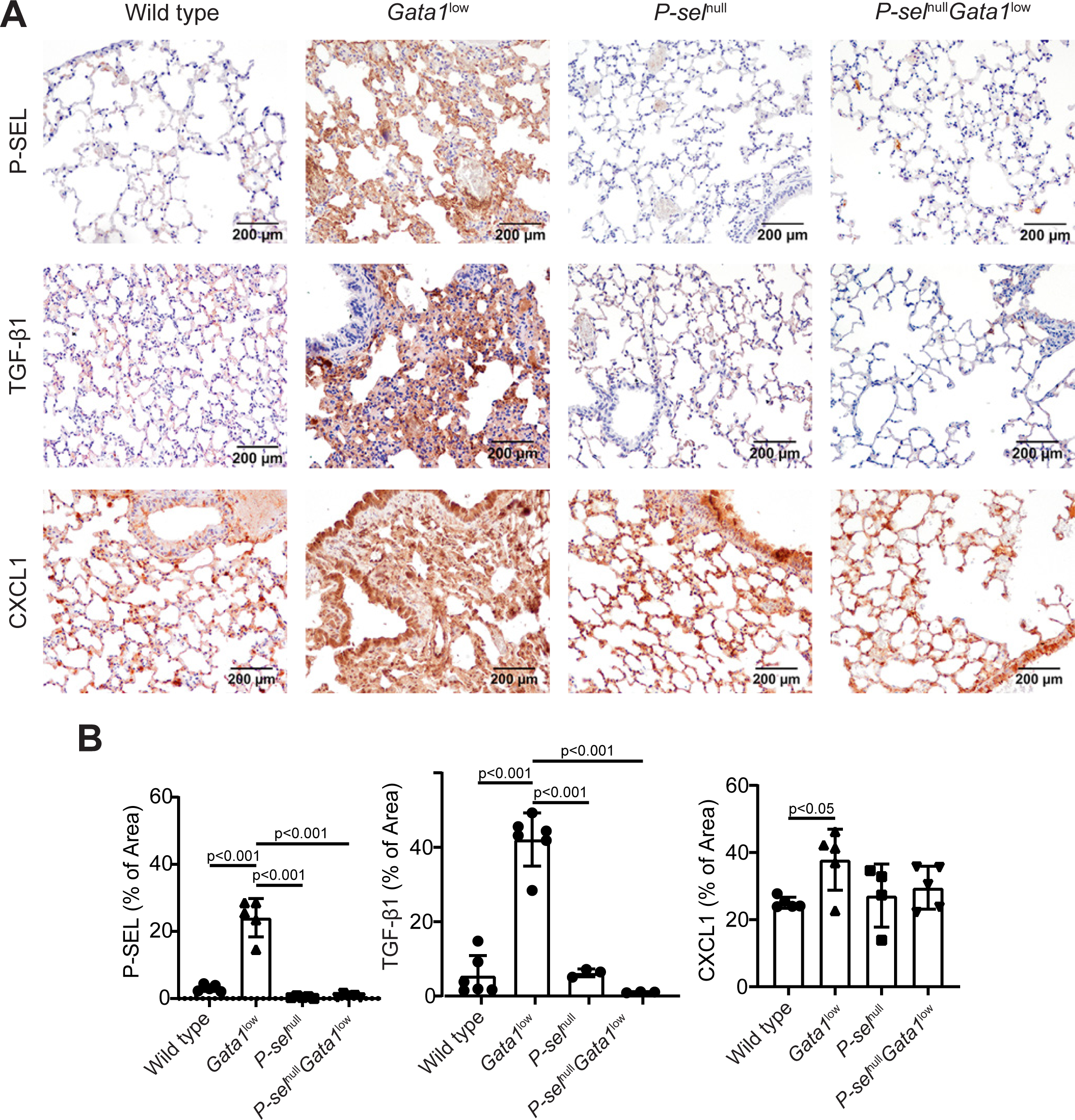
Lungs from old *Gata1^low^*mice contain levels greater than normal of P-SEL, TGF-β1 and CXCL1. A) Immunohistochemical staining for P-SEL, TGF-β1 and CXCL1 of lung sections from representative old WT, *Gata1^low^*, *P-sel^null^* and *Gata1^low^P-sel^null^* mice, as indicated. Magnification 200x. B) Quantification by computer assisted imaging of the P-SEL, TGF-β1 and CXCL1 content in the lungs from the four experimental groups. Data are presented as Mean(±SD) and were analyzed by Tukey’s multiple comparisons test. Values observed in individual mice are presented as dots. Statistically significant p values (<0.05) among groups are indicated withing the panels.

The relatively simple histological architecture of the lung allowed us to identify the cell types in which these pro-inflammatory proteins are expressed (**Figure 9**). TGF-β1 and CXCL1 were expressed at comparable levels by endothelial cells, macrophages and neutrophils from lungs of both WT and *Gata1^low^* mice while P-SEL and TGF-β1 were clearly expressed at levels greater than normal in the alveolar type II cells and in MK from *Gata1^low^* mice. The high content of P-SEL and TGF-β1 detected in *Gata1^low^*MK by immune-histochemistry is consistent with their high levels (*TGF-*β*1*: fold-change 1.49, *p*=0.006) and *P-sel*: fold change 1.63, *p*=0.031) mRNA detected by RNAseq of *Gata1*^low^ MK (**Table 2**).

**Figure 9.**
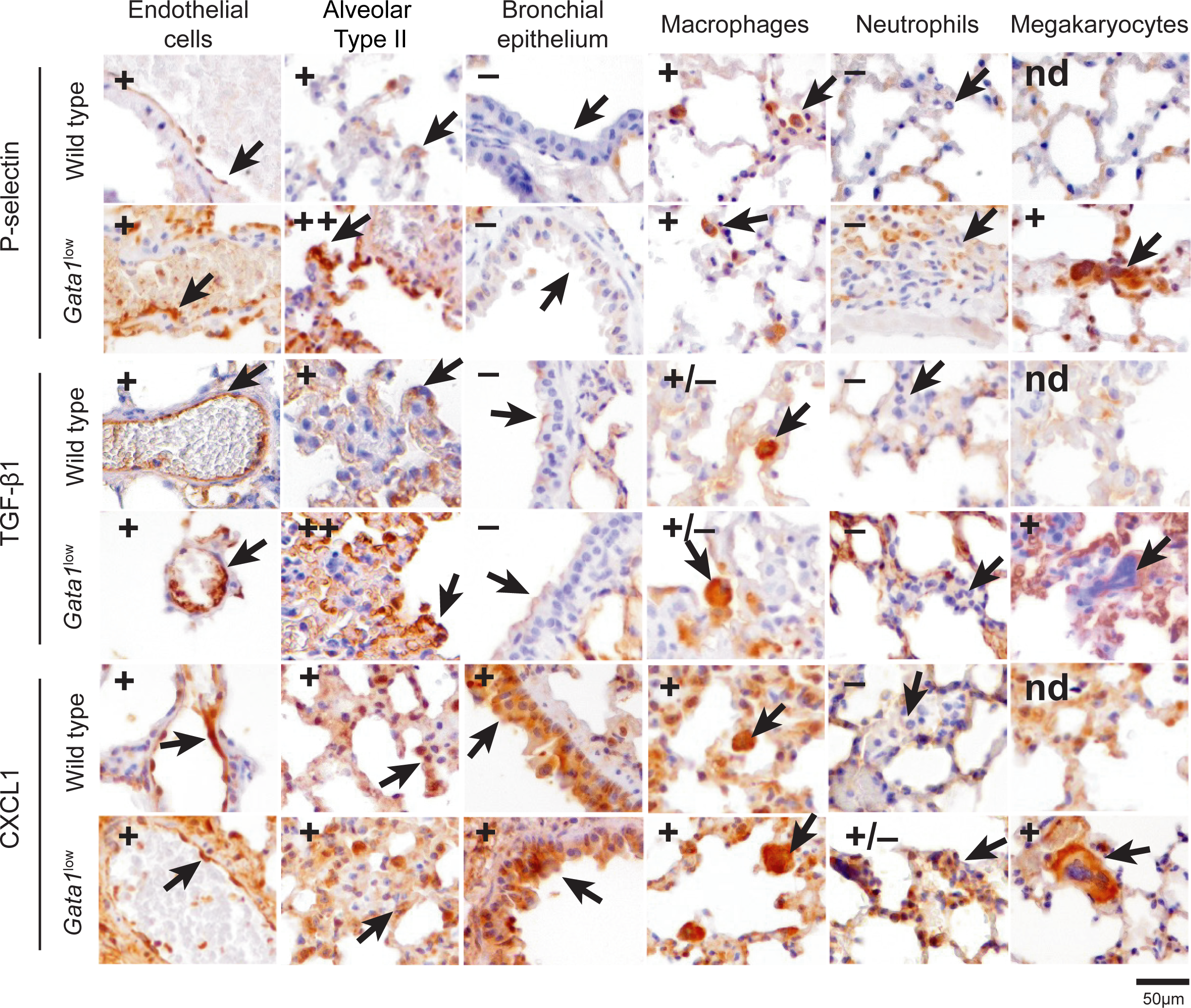
While P-SEL, TGF-β1 and CXCL1 are contained at high levels in multiple cell types in the lung from WT and *Gata1^low^* mice, they are contained at high levels in megakaryocytes only in that from *Gata1^low^* mice. P-SEL, TGF-β1 and CXCL1 content in endothelial cells, alveolar type II cells, bronchial epithelium, macrophages, neutrophils and MK from representative WT and *Gata1^low^* littermates (both 13-months old) as indicated. Cells were identified according to the following morphological criteria: Endothelial cells were recognized as flat cells delimiting the vessels lumina; alveolar type II cells were identified as cuboidal cells along the alveolar walls; bronchial epithelium was defined as pseudostratified to mono-stratified cuboidal cells along the airways of the bronchial area; macrophages were identified on the basis of their round nucleus and abundant cytoplasm; neutrophils on the basis of their small size and kidney shaped nuclei and MK on the basis of their size (10 times greater than that of any other cell type) and the polylobate morphology of their nuclei. Magnification 400x. Semi-quantitative estimates of protein content are indicated as undetectable (-), present (+) and strongly present (++).

These results suggest that high expression of P-SEL, TGF-β1 and CXCL1 by endothelial cells, bronchial epithelium and macrophages are drivers for the inflammation observed in the lung of both WT and *Gata1^low^* littermates, but that their expression by alveolar Type II cells and MK is responsible for fibrosis which is developed only by the lungs of the mutant mice.

### Fibrosis in the lung from Gata1^low^ mice is prevented by genetic ablation of P-selectin

To assess the role of P-SEL in the pathogenesis of lung fibrosis, we compared the MK profiling and the histopathology of lungs from *Gata1*^low^ mice with and without the *P-sel* gene with age (**Figures 3A**,**7C**-E, **8**, **10, S3A, S8** and **Tables 1, S2**).

**Figure 10.**
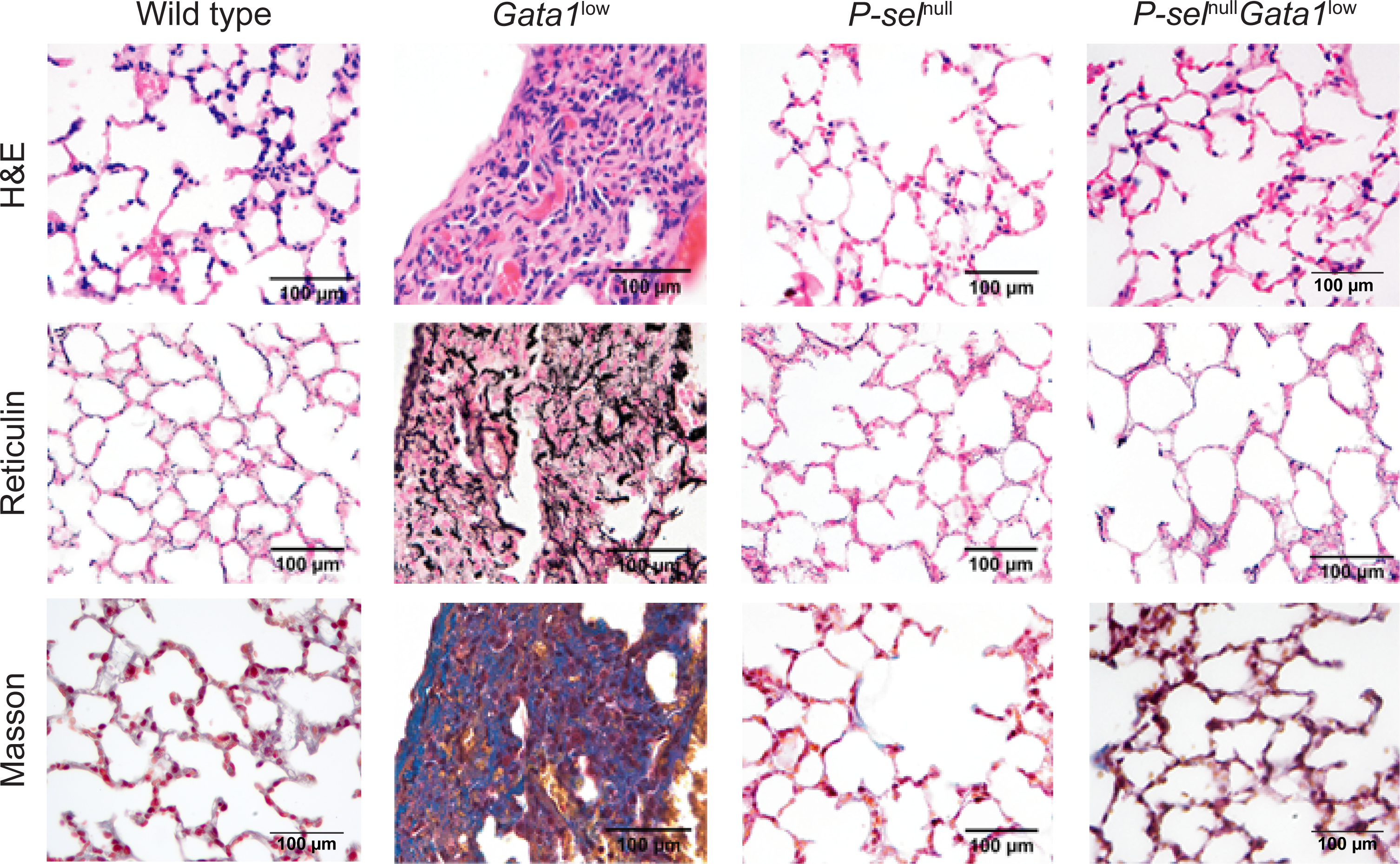
Genetic ablation of P-selectin prevents the development of fibrosis in the lung from *Gata1^low^*mice. A) H&E, Reticulin and Masson’s trichrome staining of consecutive sections of lungs from representative WT, *Gata1^low^*, *P-sel^null^* and *P-sel^null^Gata1^low^* mice (all 12-15 months old), as indicated. Magnification 40x. Results observed in multiple mice per experimental group are presented and analyzed in detail in Figure 7 and **Table 1**.

In the lung of mice carrying only the *P-sel*^null^ mutation, used as control, the total numbers of MK were similar to those observed in WT littermates (**Table S2**) although cells with a large morphology were more frequent than normal (**Figures 3A, S3A** and **Table S2).** Moreover, the lungs from *P-sel*^null^ mice manifest levels of inflammation not significantly different from that expressed by WT mice but do not express fibrosis (**Figure 10**, **7C-E** and **Table S2**). These results indicate that deletion of *P-sel* has modest effects on the baseline inflammatory phenotype of CD1 mice.

The lungs from *Gata1*^low^ and *P-sel*^null^*Gata1*^low^ mice contain similar numbers of MK (**Figure 3A**, and **Table 1**). There was a trend toward higher levels of severe (grade 2/3) inflammation in the *Gata1*^low^ lung compared to *P-sel*^null^*Gata1*^low^ mice (43.2% vs 27.8%) which is not statistically significant (unadjusted p=0.27; age-adjusted p=0.08; sex-adjusted p=0.34) (**Figure 10**, **Table 1**). However, lung fibrosis is higher among *Gata1*^low^ compared to *P-sel*^null^*Gata1*^low^ mice (Mean [±SD]=12.8 [±11.56] vs 5.2 [±5.3], unadjusted p=0.01). After adjusting for differences in sex or in age (age-adjusted p=0.04; sex-adjusted p=0.002), the difference in lung fibrosis between the two group is statistically significant (**Figure 7C-E**, **Figure 10**, **Table 1**).

Mechanistically, reduction of fibrosis in mice lacking *P-sel* is associated with barely detectable levels not only, as expected due to the genetic lesion, of P-SEL, but also of TGF-β while the levels of CXCL1 remained as high as in WT mice (**Figure 8**). As expected, single cell analyses indicate that genetic deletion of *P-sel* reduced P-SEL content in endothelial cells, alveolar Type II cells, bronchial epithelium, macrophages but did not affect the content of TGF-β and CXCL1 in these cells with respect to WT (compare **Figure S8 with Figure 9**). By contrast, deletion of *P-sel* in *Gata1^low^* mice specifically reduced the content of TGF-β in the MK with respect to that observed in *Gata1^low^* mice (compare **Figure S8** with **Figure 9**), providing support for the hypothesis that MK are the driver for the increased microenvironmental bioavailability associated with lung fibrosis in *Gata1^low^* mutants.

In conclusion, ablation of *P-sel* has modest effects on inflammation but, after adjusting for age and sex, significantly reduces fibrosis in the lungs of *Gata1*^low^ mice. Reduction of fibrosis is associated with reduced frequency of morphologically immature MK and of MK producing TGF-β1.

### Inhibitors of P-selectin, TGF-***β*** or CXCL1 rescues the fibrosis in the lung from Gata1^low^ mice

To assess the pathobiological role of pro-inflammatory factors in the development of lung fibrosis in *Gata1*^low^ mice, we tested whether it could be reversed by treating 12-months old *Gata1*^low^ mice with the P-SEL neutralizing antibody RB40.34, or the TGF-β1 trap AVID200 or the CXCR1/2, the receptors for CXCL1, inhibitor Reparixin (**Figure 11**).

**Figure 11.**
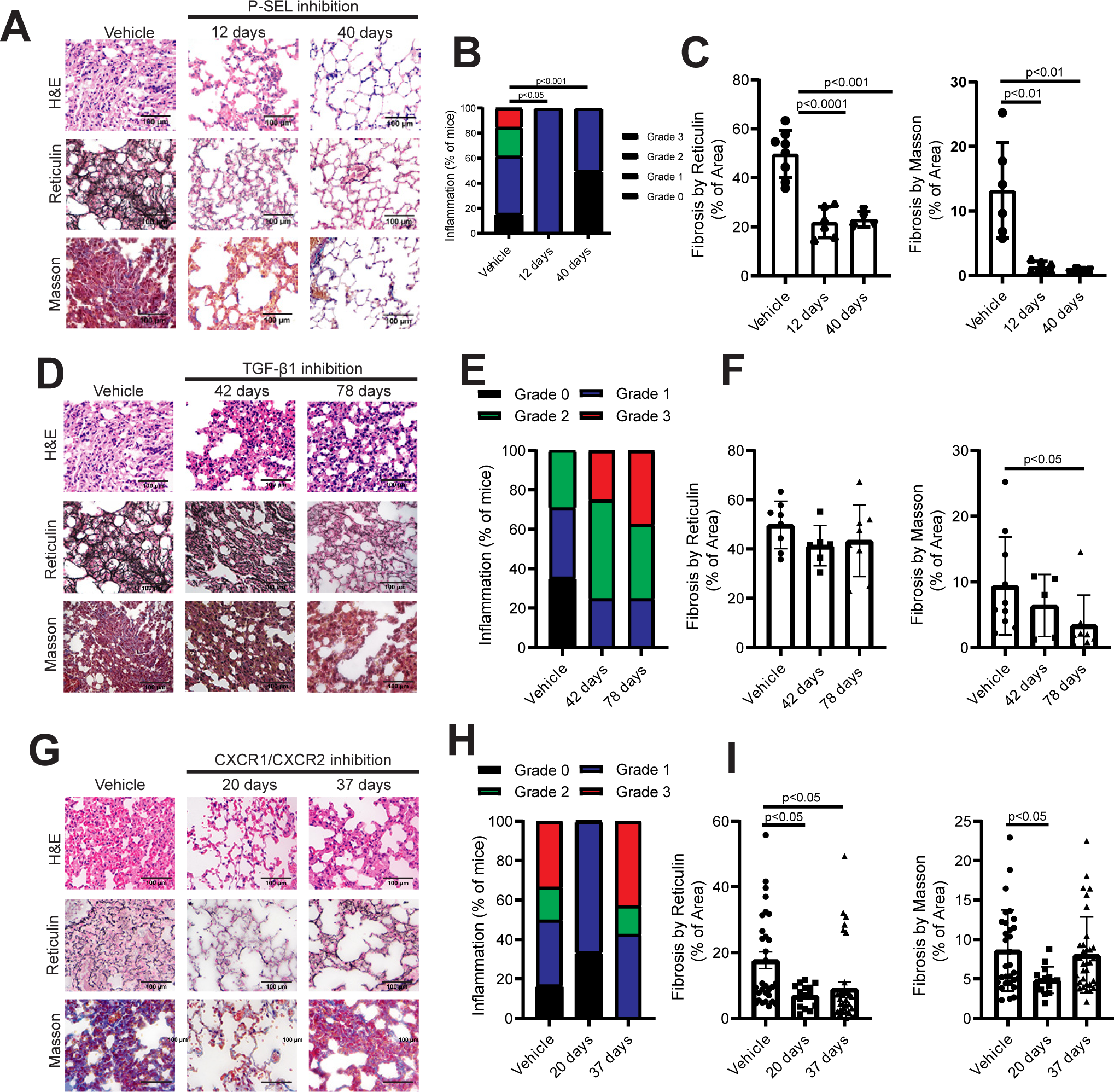
Treatments with inhibitors of either P-selectin, TGF-β or CXCR1/2 all prevent the development of fibrosis in the lung from old *Gata1^low^* mice. A, D and G) H&E, Reticulin and Masson’s trichrome staining of consecutive sections from lungs of representative *Gata1^low^* mice treated either with vehicle or the P-SEL antibody (12 or 40 days, B), or the TGF-β inhibitor (42 and 78 days, E) or the CXCR1/2 inhibitor (20 or 37 days). H), as indicated. Magnification 200x. C, F and I) Number of *Gata1^low^*mice (in percent of the total) expressing Grade 0, 1, 2 or 3 of inflammation when treated with either vehicle or the P-SEL (C), TGF-β (F) or CXCR1/2 (I) inhibitor. Results of the vehicle groups at different time points were pooled, because similar. D, G, J) Level of fibrosis detected by reticulin or Masson staining in consecutive lung sections from mice treated either with vehicle or with the P-SEL (D), the TGF-β (G), or the CXCR1/2 (I) inhibitor. Data are presented as Mean (±SD) and as data per individual mouse (each dot a single mouse). Statistical analyses were performed by one-way ANOVA and p values statistically significant between groups are indicated within the panels.

The P-SEL neutralizing antibody significantly reduced both inflammation and fibrosis in the lungs. By day 40, 50% of the treated mice had no sign of lung inflammation, and fibrosis persisted at low level by reticulin staining and was undetectable by Masson staining (**Figure 11A-C**).

The TGF-β1 trap had modest effects on lung inflammation and did not reduce fibrosis by reticulin staining although it significantly decreased the collagen fibers detected by Masson by day 78 **(Figure 11D-F**).

CXCR1/2 inhibition transiently reduced lung inflammation by day 20 and significantly reduced fibrosis by reticulin staining both at day 20 and 37 and by Masson staining at day 20 (**Figure 11G-I)**.

In conclusion, the P-SEL, TGF-β1 or CXCL1 inhibitors as single agents are all effective in reverting fibrosis in the lungs from *Gata1*^low^ mice although a dissociation with the effects on inflammation and a temporal gradient of efficacy was observed among the three drugs with inhibition of P-SEL being the most effective.

Mechanistic insights on the effects exerted by the three pro-inflammatory inhibitors on the fibrosis of the lung were obtained by investigating their effects on the TGF-β1, CXCL1 and GATA1 content of the MK (**Figures 12,13** and **S9,S10**). Treatment with the P-SEL inhibitor significantly reduced both the TGF-β1 and CXCL1 content by day 12 and the effects persisted up to day 40 **(Figure 12)**. At the single cell level, treatment with the P-SEL inhibitor reduced the TGF-β and CXCL1 content in the MK (**Figure S9**). Treatment with this inhibitor also increased the GATA1 content of the MK (**Figure 13**). By contrast, treatment with either the TGF-β1 or the CXCR1/2 inhibitor significantly reduced only the CXCL1 content by the end of the treatment without affecting that of TGF-β1 (**Figure 12**). These reductions also were more pronounced at the level of the MK (**Figure S9**). The TGF-β1 and the CXCR1/2 inhibitor did not significantly affect the GATA1 content of MK which remained low (**Figure 13, S10**).

**Figure 12.**
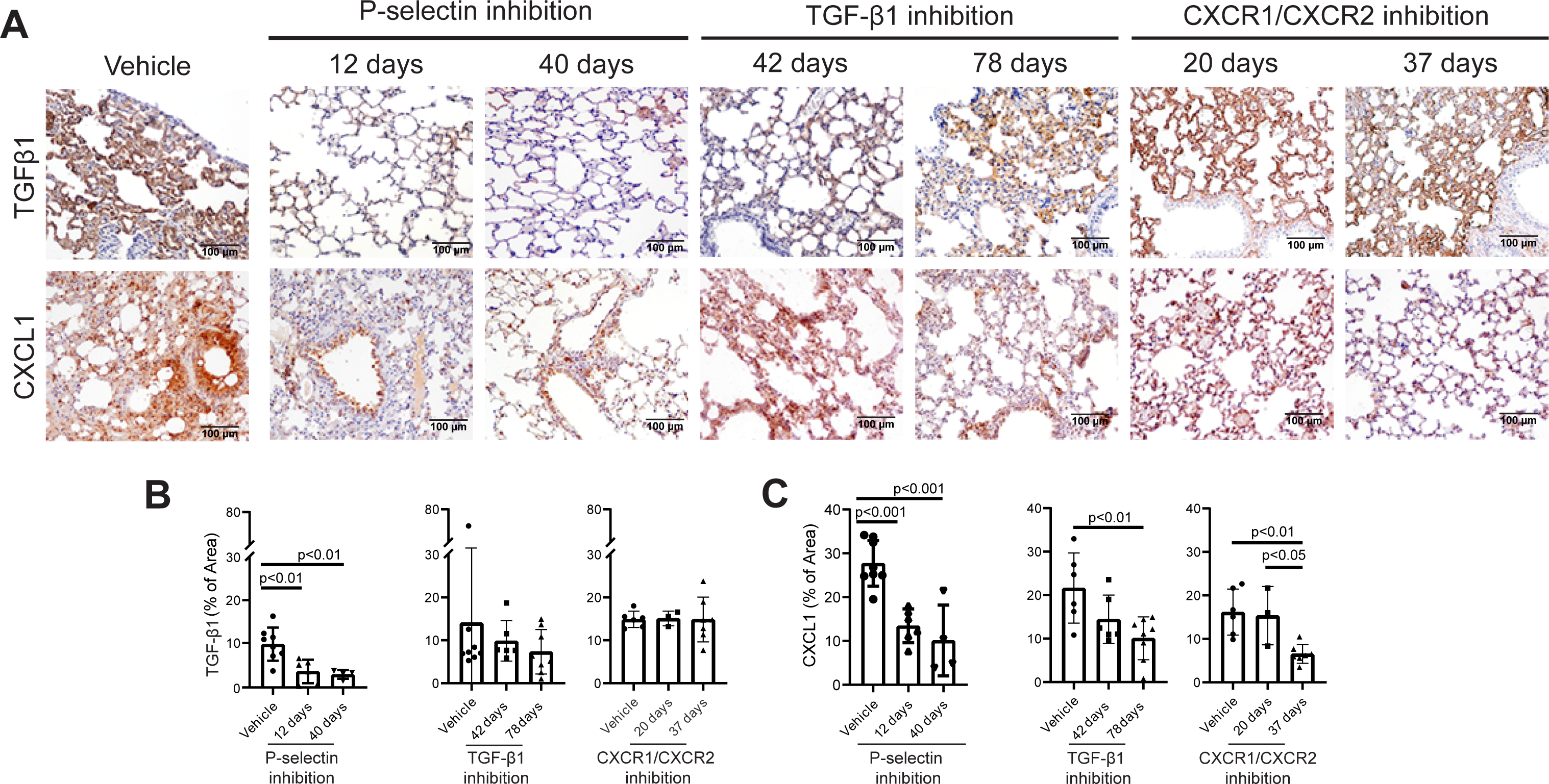
In the lung from *Gata1^low^* mice, treatment with the P-SEL inhibitor reduces the content of both TGF-β1 and CXCL1 while those with the TGF-β1 or the CXCR1/2 inhibitor reduce the levels of CXCL1 only. A) Immunohistochemical staining for TGF-β1 and CXCL1 of consecutive lung sections from representative *Gata1^low^* mice treated either with vehicle or with the P-SEL antibody (12 or 40 days), or the TGF-β (42 and 78 days) or the CXCR1/2 (20 or 37 days) inhibitor. Magnification 400x. B,C) Quantification of the TGF-β1 (B) and CXCL1 (C) content of lungs from *Gata1^low^* mice treated either with vehicle or inhibitors for P-SEL, TGF-β1 or CXCR1/2, as indicated. Data are presented as Mean (±SD) and as values per individual mice (each symbol a different mouse) and were analyzed by one-way Anova. P values statistically significant are indicated within the panels.

**Figure 13.**
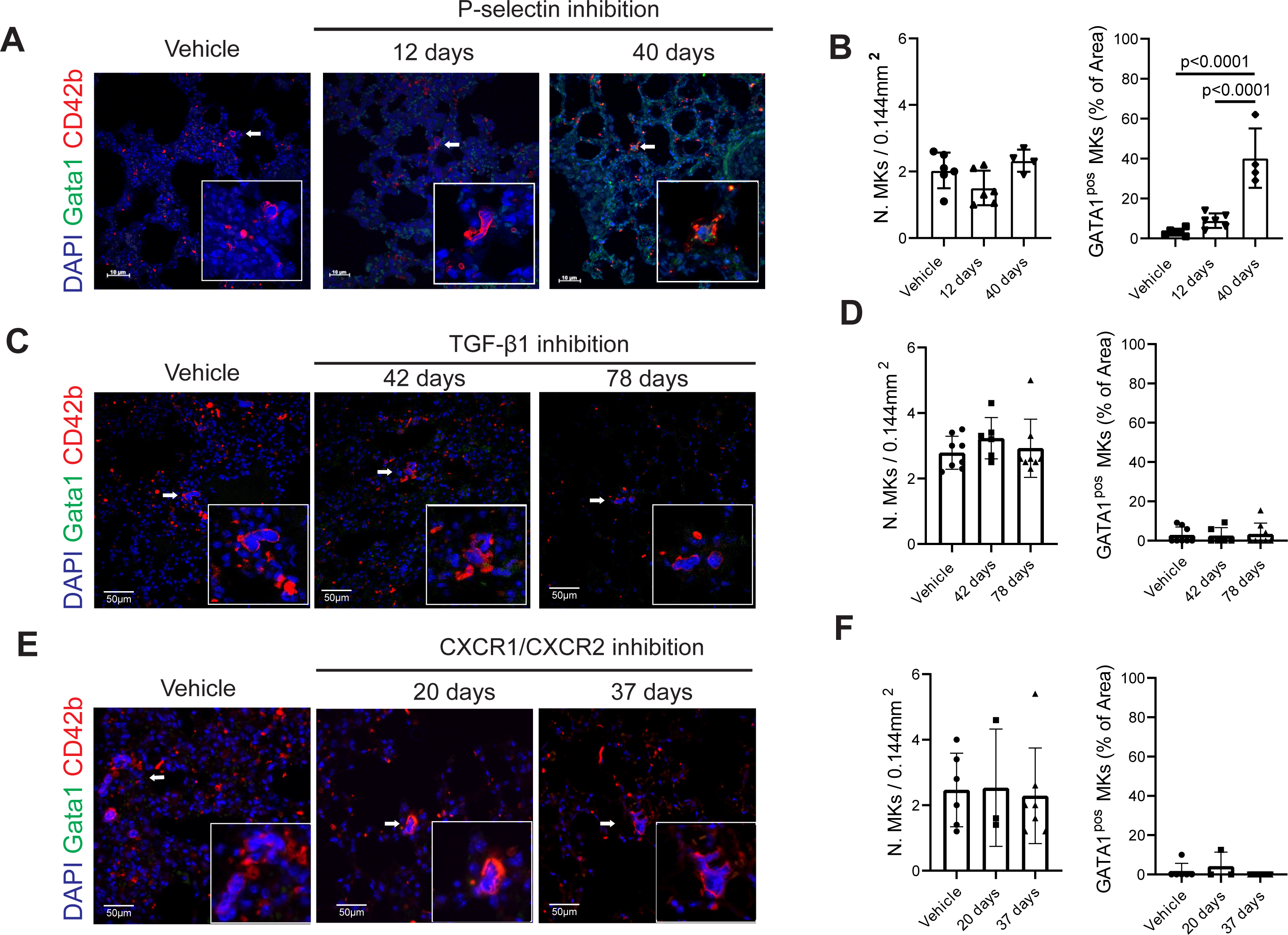
Only the treatment with the P-SEL inhibitor increases the GATA1 content of the mega-karyocytes in the lung from *Gata1^low^*mice. Double immune-fluorescence staining with anti-GATA1 (green) and CD42b of lung sections from representative *Gata1^low^* mice treated either with vehicle or with the P-SEL (day 12 and 40), TGF-β1 (42 and 78 days) or CXCLR1/2 (20 or 37 days) inhibitors, as indicated. Nuclei were highlighted with DAPI (blue). Magnification 400x. The signal captured in the individual channels are presented in Figure S10). Total number of MK (CD42bpos cells) and of MK expressing detectable levels of GATA1 in the lungs from *Gata1^low^*mice treated either with vehicle or the P-SEL, TGF-β1 or CXCR1/2 inhibitor, as indicated. Data are presented as Mean (±SD) and as values per individual mice (each symbol a different mouse) and were analyzed by one-way Anova. P values statistically significant are indicated within the panels.

These results provide further support to the hypothesis that in *Gata1^low^* mice IPF is sustained by pro-inflammatory proteins, such as TGF-β1 and CXCL1, produced by the abnormal MK.

## Discussion

Studies on IPF are challenged by the fact that most of the anatomo-pathology biorepositories contain lung specimens from a very limited number of IPF patients and from virtually no healthy individuals. We succeeded obtaining lung sections from three IPF patients and overcame the unavailability of healthy controls by analyzing sections from the lung of three patients with lung cancer which did not include evident tumor tissue. Using these specimens, we determined that although the frequency of MK was similar in the two groups, there were important differences in terms of nuclear GATA1 content (barely detectable in IPF), localization (within blood vessels in No-IPF and in the parenchyma near the alveolar epithelium in IPF) and subpopulations (enriched for immune-MK in IPF) between the MK in the two groups. These results suggest that immune-MK corrupted by low GATA1 content exert a pathological role in the etiology of IPF.

We then validated these observations by determining whether the lung from mice harboring the *Gata1^low^* mutation which impairs expression of this gene only in cells of the hematopoietic lineage and more specifically in MK, contain MK biased toward the immune lineage and develop an IPF-like phenotype with age. Of note Gata1 is not expressed in the non-hematopoietic cells within the lung. We describe that the lungs from old *Gata1^low^* mice contain MK expressing low levels of GATA1, an immature morphology (paucity of Plt territories and low ploidy), biased toward the immune lineage and localized mostly in the parenchyma near the alveoli. Furthermore, *Gata1^low^* mice develop with age fibrosis in the lung. These results support the hypothesis suggested by the human studies described above that abnormalities in the immune-MK populations sustained by low GATA1 content represent a driver for IPF.

Overall, the histological features of the fibrosis observed in the lung from *Gata1^low^* mice are more similar to that observed in IPF in patients than that displayed by the currently available bleomycin model of the disease: i) as IPF patients, *Gata1^low^* mice progressively develop both reticulin and collagen fibrosis in the lungs as they age, recapitulating the chronic nature of the disease, while in the bleomycin model fibrosis is an acute event retained until the insult persist; ii) although the lung from *Gata1^low^* mice never acquires the honeycomb morphology typical of those from IPF patients, in the *Gata1*^low^ lung, as in those from the patients, fibrosis is present predominantly in the subpleural region of the basal lobes and, occasionally, in the para-septal parenchyma while in bleomycin-treated animals, fibrosis has a bronco centric localization that reflects the site where the insult is applied; iii) as in some IPF patients, inflammation is not the primary cause of fibrosis in the lung from *Gata1^low^* mice while fibrosis is a direct consequence of inflammation in bleomycin-treated animals. Our conclusion that inflammation is not the primary driver of fibrosis in *Gata1^low^* mice is based on robust considerations: i) inflammation is observed also in the lung from WT and *P-sel^null^* mice which never develop fibrosis; ii) genetic deletion of *P-sel* prevents development of fibrosis in the lung of mice carrying the *Gata1^low^* mutation but does not reduce the base-line inflammation of this organ; iii) pharmacological inhibition of either P-SEL, TGF-β or CXCR1/2, has modest, if any, effect on inflammation but rescues the fibrosis in the lung of *Gata1^low^* mice. We suggest that inflammation is a confounding factor rather than a cause of fibrosis in IPF and that *Gata1^low^* mice represent a model for IPF more stringent than the bleomycin-treated mice currently used to study the etiology of this disease. Additional studies comparing the megakaryocyte subpopulations present in the lungs from IPF patients and from patients with inflammation but not fibrosis, such as idiopathic non-specific interstitial pneumonia, chronic hypersensitivity pneumonitis or connective tissue disease related idiopathic lung disease, are necessary to clarify the MK abnormalities that specifically drive IPF in patients, This study will be performed in the near future under the umbrella of PUMA, a retrospective observational protocol approved by the Human Subject Committee of Campus Biomedico and Policlinico Gemelli on March 21^st^, 2023.

It is very interesting that the anatomical localization, and other features, of the fibrosis in the lung of *Gata1^low^* mice are very similar to that of the recently described Tamoxifen-treated *SP-CI73T* mouse model^25^. Furthermore, in the lung of *Gata1^low^* mice the corrupted MK are found closely associated with the alveolar epithelium and are surrounded by fibers of collagen 1A and III. These observations induce us to speculate that *Gata1^low^* MK sustain fibrosis by secreting factors that alter the metabolism of the surfactant proteins expressed by the alveolar type II cells turning them into pro-fibrotic cells. Additional support for this hypothesis is that the content of proinflammatory cytokines of the alveolar Type II cells in *Gata1^low^*lung, as those of the corresponding cells from Tamoxifen-treated *SP-CI73T* mouse model^25^, include high levels of TGF-β.

In the bleomycin-animal model, inflammation is initially associated with accumulation of neutrophils within the lung, which are subsequently replaced by lymphocytes^60^. This accumulation is associated with increases in the levels of systemic and microenvironmental pro-inflammatory cytokines (TGF-β, CXCL1 and others), which precede the appearance of the pro-fibrotic markers (fibronectin and pro-collagen 1) and of fibrosis^5^. Abnormal expression of P-SEL by endothelial cells^10^ and of TGF-β by numerous cell types, including endothelial cells, epithelial cells, alveolar type II cells and macrophages^53, 54^, has been implicated in the development of fibrosis in the lungs of bleomycin-treated animals. High levels of TGF-β produced by the alveolar Type II cells were also associated with the development of fibrosis in the Tamoxifen-treated *SP-CI73T* mouse model^25^. By contrast, although CXCL1 is elevated in bronco-alveolar lavages of these mice^61^, the cell population responsible for its production in the lung of mouse models has yet to be identified. Most of the findings in mice have been validated by studies involving IPF patients^55, 57, 58, 62 50–52^. An important exception is P-SEL that in the patients is expressed at high levels not by the endothelial cells but by Plt, cells which originate from the MK^49^. The debate on which cell(s) is responsible to establish the proinflammatory molecules milieu responsible for fibrosis in the lung is expected since the studies in all the models available up to now, including the Tamoxifen-treated *SP-CI73T* mice, are confounded by inflammation. The fact that the strain hosting the *Gata1^low^*mutation is well known to express a baseline inflammatory state with age, allowed us to use a combination of genetic and pharmacological approaches to dissect the identity of the cells specifically responsible to sustain fibrosis in the lung of our model.

By immune-histochemistry, endothelial and epithelial cells, alveolar macrophages and interstitial fibroblasts from the lung of WT and *P-sel^null^*mice, which express inflammation but no fibrosis, contain robust levels of P-SEL, TGF-β and/or CXCL1, suggesting that the pro-inflammatory profile of these cells is responsible for the baseline inflammation of the CD1 strain. By contrast, MK, and to some extent alveolar type II cells, are the only cells that contain robust levels of these pro-inflammatory mediators specifically in the lung from *Gata1^low^* mice, suggesting that they are responsible to establish the pro-inflammatory milieu that drives fibrosis in our model. Support for the pro-inflammatory role of the MK is provided by their RNA-seq profiling indicating that those of the lung from *Gata1^low^* mice express upregulated levels of TGF-β (fold-change=1.49, *p*=0.006) and CXCR2, the receptor for CXCL1^63, 64^ (fold change=1.95, p=0.007) while the levels of CXCL1 were downregulated (fold-change=-2.843, P=0.000). The discrepancy between the reduced levels of *CXCL1* mRNA detected in *Gata1*^low^ MK by RNA-seq and their high *CXCL1* content detected by immunohistochemistry may be explained by the high levels of mRNA for *CXCR2*, a receptor which internalize its ligand without routing it to the degradation system, expressed by these cells which allow for greater accumulation of the protein in the cytoplasm. The RNAseq profiling also displays downregulation of other factors involved in the MK-specific anti-microbial response in this organ: *CXCL1* fold-change=2.8, *p*=0.000), *CXCL5* (fold-change - 2.691, *p*=0.017, and *CXCL10* (fold-change -1.88, *p*=0.033)^63–66^, confirming that the GATA1-deficient immune MK detected in the lung from old *Gata1^low^* mice are corrupted.

The pathogenetic role of P-SEL, TGF-β and CXCL1 in the fibrosis of the lung from *Gata1^low^* mice was further dissected by evaluating the effects of the deletion of *P-sel* and pharmacological inhibition of P-SEL, TGF-β and CXCR1/2 on the lung phenotype of these mice. Genetic deletion of *P-sel* prevented mice carrying the *Gata1*^low^ mutation to develop fibrosis in the lung with age. This effect was associated with reduced content of both TGF-β and CXCL1 in the MK from this organ. Furthermore, inhibition of P-SEL reduced fibrosis (both as reticulin and collagen fibers), the frequency of MK with low GATA1 content and the content of TGF-β and CXCL1 in the lung for up to day 40 of treatment. By contrast, TGF-β inhibition reduced collagen fibers and CXCL1 content only by the end of the treatment (day 72) while that of CXCR1/2 reduced collagen fibers transiently (day 20 only), reticulin fibers both at day 20 and 37 and CXCL1 content only at day 37. The efficacy of the CXCR1/2 inhibitor to consistently reduce only reticulin fibers is consistent with the reported efficacy of this drug in Bleomycin-treated mice that are a model for early stages of the disease^67, 68^. Of note, both the TGF-β and CXCR1/2 inhibitors did not affect neither the TGF-β nor the GATA1 content of the MK, which remained, respectively high and low, in the lung of the treated *Gata1*^low^ mice. Although the effects exerted by the potent and specific TGF-β1 trap on the fibrosis of the lung from *Gata1*^low^ mice are inferior to those reported for additional drugs which target the same pathway tested in bleomycin-treated models, they are consistent with the poor performance of those clinically tested up to now in IPF^14–17^.

Overall, these data suggest that in *Gata1*^low^ mice fibrosis in the lung is triggered by an abnormal population of immune-MK which initiates a sequence of pathological effects exerted by multiple pro-inflammatory proteins over time. It may be speculated that the first step is accumulation of immune-MK that became engaged into a process of pathological emperipolesis mediated by high levels of P-SEL^28^. This pathological emperipolesis leads to death of the MK by para-apoptosis and release of the content of their cytoplasm into the microenvironment, increasing the bioavailability of TGF-β and CXCL1, and possibly of other proinflammatory cytokines not identified as yet, in the microenvironment. The release of these pro-inflammatory cytokines sustains a mesenchymal cell transition responsible for fibrosis. Since the *Gata1*^low^ MK from the lung, although embedded in collagen fibers, do not contain these proteins in their cytoplasm, we exclude fibrosis is mediated by niche-poised MK, a population responsible to synthesize the extracellular matrix during organo-genesis^45^ and that, when pathologically reactivated in adulthood, promote scarring of the BM^28^. We suggest that, as in Tamoxifen-treated *SP-CI73T* mice, the cells undergoing the mesenchymal transition responsible for fibrosis in the *Gata1*^low^ lung are the alveolar Type II cells which became corrupted by the nearby MK.

In addition to the poor fidelity of the No-IPF controls used in the human experiments discussed above, our study has other limitations: i) the mutation status of the IPF patients analyzed is not known. However, given the low frequency of the telomerase-related or surfactant proteins mutations found in IPF patients, it is unlikely that our patients harbored any of the mutations already known; ii) the *Gata1^low^* mutation reduces the content of the protein in all the megakaryocyte subpopulations and not only in those with immune functions. Unfortunately, conditional knock-out mice with reduced/ablated GATA1 expression only in the immune MK are not available. The available conditional *PF4 Cre* strain^69^ is not suited for our studies because in this strain GATA1 expression is completely ablated in all the megakaryocyte sub-populations, making impossible to identify abnormalities induced by reduced GATA1 content in specific MK subpopulations. Since, compared to the other subpopulations, the development of immune-MK is specifically sensitive to the expression of *Pu1*, double mutants in which the *Gata1^low^* mutation is combined with conditional *PF4 Cre-Pu1* knock-out mice may be more suited to rigorously assess the role of immune MK in the development of IPF. Unfortunately, these mutants are not yet available; iii) our assumption that the corrupted *Gata1^low^* MK and the altered alveolar Type II cells are casually linked is based only on the anatomical localization of the two cell types. Further studies, which will rigorously assess whether the *Gata1^low^*MK alter the functions of alveolar Type II cells by determining their expression profiling using specific lineage-tracking genetic mouse models, which again are not currently available, are necessary to formally assess whether MK and alveolar type II cells are related in the development of IPF of our model.

In spite of these limitations, we believe that our data are important because they open new perspectives for the etiology and cure of a disease the genetic lesion of which is currently unknown for >95% of the patients and that has limited therapeutic options. In this regard, cells of BM origin had been already associated with development of IPF^70^, leading to the interesting hypothesis that, at least in some patients, IPF may have a hematopoietic origin that may be treated with BM transplantation^71^. However, the consideration that IPF patients are old and fragile and unlikely to survive the fully ablating conditioning regimes available in 2004 when these studies were performed has left this hypothesis unexplored. The more patient friendly regimens under development (i.e. hematopoietic stem cell depletion of the host by treatment with anti-cKIT antibodies)^72–74^ may make now this approach feasible even in IPF patients. The fact that reduced conditioning in transplanted patients with myelofibrosis, a population as fragile as that affected by IPF, gave satisfactory results supports the feasibility of transplants with this regiment in older patients^75^. This hypothesis deserved to be tested by proof-of-principle experiments in animal models.

In conclusion, *Gata1^low^* mice are a novel genetically driven IPF animal model and provide strong evidence that immune-MK alterations play a major role in the development of this disease. The *Gata1*^low^ model also clarifies that P-SEL dysregulation predates TGF-β and CXCL1 in the pathobiology of IPF and should be considered for treatment of these patients.

## Materials and Methods

### Human Subjects

Consecutive lung sections from IPF and non-IPF patients (three different patients per group) were provided, respectively, by the Institute of Human Pathology, Policlinico Sant’Orsola, Bologna, Italy and by Policlinico Campus Bio-Medico (Rome). Details of the source of the subjects included in the study are provided in **Table S1**.

### Mice

Male and female WT, *P*[*sel*^null^, *Gata1*^low^ and *P*[*sel*^null^*Gata1*^low^ mice were generated in the animal facility of Istituto Superiore di Sanità which is the recipient of the NIH animal welfare assurance number F16-00089, as previously described^40^. All the mutations are in the CD1 background. The study was approved by the Italian Ministry of Health (protocol n. 419/2015-PR and D9997.121) and conducted according to the European Directive 2010/63/EU for the protection of animals used for scientific purposes.

### Treatments

*<u>P-SEL inhibitors:</u>* Twenty 8-11 months-old *Gata1*^low^ mice were randomly divided in two groups which were treated either with vehicle (2% v/v dimethyl sulfoxide in H_2_O) or with biotinylated mAb RB40.34 (BD Pharmigen, San Diego, CA; 30 μg/mouse every other day by tail vein injection), as described^76, 77^. Mice were sacrificed after 12-40 days for histopathologic determinations. *<u>TGF-</u>*<u>β</u>*<u>1 inhibitors</u>*: *Gata1*^low^ mice (10 - 12 months-old) were weighted and randomly divided into two groups (four males and four females/group) and treated with either vehicle (PBS) or with the TGF-β1 trap AVID200 (5mg/Kg twice a week by intraperitoneal injection), as described^78^. Mice were treated either for 42 (six mice, three females and three males with AVID200 and 2 females with vehicle) or 72-78 (eight mice, four females and four males in each group) days. At day 42, 72 or 78 from the beginning of the treatment, mice were euthanized, and their lung removed for histopathology observations. *<u>CXCR1/CXCR2 inhibitors</u>*: Sixteen eight-month-old *Gata1*^low^ mice were anesthetized with 2-to-3% isoflurane and implanted subcutaneously with an ALZET^®^ Osmotic Pump (model 2002) pre-filled with 200μL of vehicle (sterile saline) or the CXCR1/CXCR2 inhibitor Reparixin (7.5mg/h/Kg in sterile saline) as described^79, 80^. Mice were sacrificed 20 or 37 days after the beginning of the treatment, and their lungs removed for histopathological determinations.

### Histochemistry

Lung and BM from WT, *Gata1*^low^, *P*[*sel*^null^ and *P*[*sel*^null^*Gata1*^low^ mice harvested from mice that had previously received intratracheal inoculation of 10% (vol/vol) phosphate[buffered formalin and then fixed in 10% (vol/vol) phosphate[buffered formalin, paraffin embedded and cut into consecutive 3μm sections, according to standard procedures. Consecutive sections from the mouse organs, as well as from BM and lungs of the patients, were stained with Hematoxylin[Eosin (H&E, cat no. 01HEMH2500 and 01EOY101000, respectively; Histo-Line Laboratories, Milan, Italy), Reticulin (Bio-Optica, Milan, Italy) or Masson’s trichrome (Bio-Optica) and evaluated with a Nikon Eclipse E600 microscope (Nikon Instruments Europe BV, Amsterdam, NL) equipped with the Imaging Source "33" Series USB 3.0 Camera (DFK 33UX264; Bremen, DE). Inflammation was assessed on de-identified H&E stained sections by an experienced veterinarian pathologist and expressed on the basis of a semiquantitative scale based on the numbers of inflammatory cells (neutrophils, macrophages and lymphocytes) that were infiltrating the alveolar septae and/or peribronchial-vascular spaces: grade 0 (absence of inflammatory cells); grade 1 (rare inflammatory cells in <10% of the area); grade 2 (moderate numbers of inflammatory cells in 10-30% of the area) and grade 3 (presence of large numbers inflammatory cells in >30% of the area). <u>Fibrosis</u>.

Semiquantitative and quantitative assessment of fibrosis were performed on sections stained with Reticulin or Masson’s trichrome staining by an experienced veterinary pathologist and by Image analysis using the software ImageJ (version 1.52t, National Institutes of Health, Bethesda, MD, USA), respectively. The semiquantitative scoring was the following: grade 0 (presence of collagen fibers only at the bronchovascular level and absence of septal alveolar fibrosis); grade 1 (presence of focal mature collagen fibers in the alveolar-septal interstitium); grade 2 (deposition of multifocal to coalescing collagen fibers in the alveolar-septa interstitium); grade 3 (deposition of collagen bundles that irregularly thicken the alveolar septa and alterations of the overall pulmonary architecture) (**Figure S6A**). Quantitative image analysis was performed on five different photomicrographs randomly selected per each lung section/mouse (area of each photomicrograph=1.49mm^2^) (**Figure S6B**). The number of mice analyzed per experimental group is indicated in the legends. Fibrosis was expressed as percent of the area stained in blue with respect to the total area of the field censored of the empty alveolar spaces. Statistical analyses confirmed a good correlation between the semiquantitative and quantitative determinations of fibrosis (**Figure S6C**).

### Immunofluorescence (IF) studies

Mouse and human lung and sections (3μm thick each) were dewaxed in xylene and treated with EDTA (buffer pH=8), incubated for 20min in a pressure cooker (110-120°C, high pressure) for antigen retrieval and then with antibodies against CD42b (cat no. ab183345, Abcam, Cambridge, UK) or/and GATA1 (cat no. sc-265, Santa Cruz Biotechnology, Santa Cruz, CA, USA), Collagen I (cat no. NB600-408, Novus biologicals, Centennial, CO, USA), Collagen III (cat no. ab7778, Abcam) or Collagen IV (cat no. ab6586, Abcam) over night at 4°C. Cell binding of the primary antibodies was visualized with a secondary antibody goat anti-rat Alexa Fluor 488 (cat no. ab150165, Abcam) or anti-rabbit Alexa Fluor 555 (cat no. ab150078, Abcam), as appropriate. Sections were counterstain with DAPI (cat no. D9542-5MG, Sigma Aldrich, Saint Louis, MI, USA) and mounted with Fluor-shield histology mounting medium (cat no. F6182-10MG, Sigma Aldrich). The slides were examined using a Nikon Eclipse Ni microscope equipped with the appropriate filter cubes to distinguish the fluorochromes employed. The images were recorded with a Nikon DS-Qi1Nc digital camera and NIS Elements software BR 4.20.01 and selected sections were acquired using FV1000 confocal microscope (Olympus, Tokyo, Japan) equipped with a confocal spectral imaging system (Olympus Fluoview 1000). The number of MK was counted in 10 fields per section and expressed as mean value.

### Confocal Microscopy

Femurs were fixed in formaldehyde (10% v/v with neutral buffer), treated for 1h with a decalcifying kit (Osteodec; Bio-Optica, Milan, Italy) and included in paraffin. Three micron-thick BM sections were dewaxed in xylene and antigens were retrieved by treatment with EDTA buffer (pH=8) for 20’ in a pressure cooker (110-120°C, high pressure). For each sample almost 3 consecutive sections were obtained. One was incubated with antibodies against CD41 (rat anti mouse, Santacruz Biotechnology INC. (MWReg30): sc-19963) and LSP1 (EPR5997), another one with antibodies against CD61 (B-7, mouse anti mouse, Santacruz Biotechnology INC., sc-46655) conjugated with Alexa 488, and MYLK (polyclonal rabbit anti-mouse, Proteintech, 24309-1-AP); and the last one incubated with antibodies against CD42b (mouse monoclonal, Santacruz Biotechnology INC, sc-271171) and ARNTL (rabbit anti mouse/human NB100-2288, Bio-Techne), for 2 hours at room temperature. After extensive washing, sections were incubated for 45 min at room temperature with the following secondary antibodies: CD41 was visualized with the antibody goat anti rat Alexa Fluor 488 (Catalog #A-11006, Invitrogen); the LSP1, CD42b and MYLK antibodies were visualized with the antibody donkey anti-rabbit Alexa Fluor 555 (Catalog # A-31572, Invitrogen) and ARNTL was visualized with the antibody goat anti mouse Alexa Fluor 488 (Catalog #A32723, Invitrogen), CD61 is directly conjugated with Alexa 488. All sections were mounted with Mounting Medium With DAPI - Aqueous, Fluoroshield (Catalog #ab104139, Abcam) and examined using a confocal microscope Zeiss LSM 900 (Carl Zeiss GmbH, Jena, Germany) in Airyscan mode, using a (Zeiss) planapo objective 60x oil A.N. 1,42. Excitation light was obtained by a Laser Dapi 408 nm (DAPI), an Argon Ion Laser (488 nm) for Alexa 488 and a Diode Laser HeNe (561 nm) for Alexa 555. DAPI emission was recorded from 415 to 485 nm, Alexa 488 emission was recorded from 495 to 550 nm, Alexa 555 emission was recorded from 583 to 628 nm. Images recorded have an optical thickness of 0.3 μm and were processed with the Zen Blue (3.2) (Carl Zeiss GmbH, Jena,Germany) and ImageJ (1.53t – National Institutes of Health, USA, http://imagej.nih.gov/ij) softwares. All images were collected with the same detector value for DAPI, Alexa 488 and Alexa 555, respectively. The background of each signal was calculated splitting the channels and the single channel background obtained in areas external to cells. The number of positive cells were obtained in the whole section(s).

### Immunohistochemistry (IHC)

Three micron-thick lung sections were dewaxed and rehydrated. Endogenous peroxidase was blocked by immersion in H_2_O_2_ in methanol (3%, v/v) for 30’ at room temperature. Antigen retrieval was performed by incubation in citrate buffer, pH=6.0 for 1min in a microwave oven at 750W, followed by cooling at room temperature for 20’. After blocking of non-specific antigenic sites by incubation with goat serum in PBS (10%, v/V) for 30’ at room temperature, slides were incubated with CD42b (cat no. ab183345, Abcam, Cambridge, UK), biotinylated anti-P-SEL (cat no. 553743, BD Pharmigen), anti-TGF-β1 (cat no. sc-130348, Santa Cruz Biotechnology), anti-CXCL1 (cat no. ab86436, Abcam), anti-CXCR1 (cat no. GTX100389, Genetex, Irvine, CA, USA) and anti-CXCR2 (cat no. ab14935, Abcam) antibodies overnight at 4°C. Binding sites were revealed with the commercial avidin-biotin-peroxidase kit (ABC Kit Elite, Vector, Burlingame, CA) and the chromogen 3,3′-diaminobenzidine (0.05%, Histo-Line Laboratories) either directly or previous incubation with suited secondary biotinylated antibodies. Slides were counterstained with Harris hematoxylin (Histo-Line Laboratories) and mounted with HistoMount DPX medium (cat no. 01BM500H, Histo-Line Laboratories). Image analysis was performed using ImageJ program and the content of antibody binding was calculated as intensity of the brown staining above the threshold in areas (1.49mm^2^ each) from 5 different photomicrographs randomly selected for each sample. The intensity of the staining of individual cells recognized according to standard morphological criteria was arbitrarily assessed as “–“ (below detection), “+” (detectable) and “++” (strongly detectable).

### Flow cytometry of lung megakaryocytes

Lungs were incubated in Dulbecco’s modified Eagle Medium (cat no. D8537, Sigma-Aldrich, Saint Lois, M0, USA), supplemented with 10% (v/v) fetal bovine serum (cat no. F7524, Signa Aldrich), penicillin/streptomycin (cat. no. 15140122), vitamins (cat no. 11120052), Glutamax (cat. no. 10566016), non-essential amino acids (cat no. 11140050) and sodium pyruvate (cat no. 11360070) plus collagenase type II (1mg/mL, cat no. 17101015) (all from Gibco; Grand Island, NY, USA), as described^46^. After 30min at 37°C, lungs were mashed through a 100-μm cell strainer (BD Pharmingen) and washed with cold media and centrifuged at 1200 rpm for 5 minutes. ACK Lysing Buffer (cat no. A1049201, Gibco) was added to the pellet for 5 minutes in ice. The ACK was washed out with 50 mL of PBS and centrifuged at 1200 rpm for 5min. Cells were then incubated with phycoerythrin (PE)-CD41 (cat no. 558040) and fluorescein isothiocyanate (FITC)-CD61 (cat. no. 553346). Dead cells were excluded by Sytox Blue staining (0.002mM, Molecular Probes, Eugene, OR, USA). For DNA content determinations, cells were then incubated with phycoerythrin (PE)-CD42b (cat. no. 555473) and FITC-CD61, washed and fixed with 4% paraformaldehyde (Sigma-Aldrich) for 60min, permeabilized with Triton-X100 (0.2% v/v in PBS, cat. no. 108643; Merck, Darmstadt, Germany) for 15min and incubated with propidium iodide (40 μg/mL in PBS) and RNase A (100units/mL, cat. no. 10109142001, Merck) for 30min at 37[. All the antibodies are from BD Pharmingen. Cells were analyzed with Gallios (Beckman Coulter, Brea, CA, USA) and the flow cytometry data processed with the Kaluza program (version 2.1, Beckman Coulter).

### RNA seq analysis

MK were purified from the BM and lungs of two *Gata1^low^* and three WT littermates (11-25 months of age. Total RNA was collected using the Direct-zol^TM^ RNA MiniPrep (>200,000 sorted MK) or MicroPrep (<200,000 sorted MK) (Zymo Reaserch). Illumina libraries were constructed using SMARTer Stranded Total RNA-Seq Kit v3 - Pico Input Mammalian (TaKaRa, Japan). RNA-seq libraries were sequenced at read lengths 2 × 151bp on the Illumina Novaseq 6000 (Illumina). Aligment was performed using RNA-STAR (2.3.1) using the ENCODE 4 default transcriptome annotation (GENCODE v25). Gene counts were obtained using featureCounts and FPKM per gene were calculated using Cufflinks. Differentially expressed genes (DEGs) between the two groups of lungs MK from *Gata1^low^* mice and WT mice were analyzed with R package DESeq2 and the GO Enrichment Analysis was performed with R package enrichplot, DOSE^81^ and clusterProfiler ^82, 83^. Multiple testing correction was performed using the Benjamini-Hochberg method and significance was established at an FDR of 5%.

### Gene Set Enrichment Analysis

GSEA was performed using GSEA software GSEA 4.3.1 wiTH default parameters. The CURATED gene set (m2.all.v2022.1.Mm.symbols.gmt) used for the analysis was downloaded from MSigDB (https://www.broadinstitute.org/gsea/msigdb/collections.jsp). A total of 32,527 expressed genes were included in the analysis. Genes were pre-ranked in order of their differential expression between MK from *Gata1^low^*lungs and from CD1 lungs.

### Statistical methods

Categorical variables are presented as counts and percentages by genotype; Fisher’s exact tests were used to test the univariate association between genotype and outcome. Continuous variables are presented as mean (standard deviation [SD]), median, and range). Statistical analyses was performed by parametric t-test or one-way ANOVA for multiple comparisons, as appropriate. Multiple linear regression and ordinal logistic regression were used to model relationships between genotype and outcomes, adjusted by age or by sex. Multiple linear regression was used to test the interaction between age and genotype on the outcome of percent fibrosis. Where interaction was found, simple linear regression was used to assess the relationships between fibrosis outcomes and age, stratified by genotype. Alignment between semiquantitative and quantitative levels of fibrosis was assessed using box plots. All hypothesis testing was two-sided, with p<0.05 used as the threshold for statistical significance. Analyses were conducted using SAS software (version 9.4; SAS Institute Inc., Cary, NC, USA), R (version 3.6.3), and Prism 8 (version 8.0.2; GraphPad Software, La Jolla, CA).

### Study approval

This is not a human study subjected to prior IRB approval according to the Declaration of Helsinki for Studies Involving Human Subjects. All human specimens were provided as de-identified material by public repositories.

### Data availability

Data will be deposited in publicly accessible databases upon acceptance of the manuscript.

## Author contributions

FG, MZ, PV, MF, FA, FM and GS performed experiments and analyzed data. MM and JV performed the RNA-seq analysis. CMH and ACD did the statistical analysis. AP, GS, SN, and LR reviewed the data and wrote the manuscript. ARM, designed the experiments, analyzed and reviewed the data and wrote the manuscript. All the authors read the manuscript, concurred with its content and approved the submission.

## Acknowledgments

This study was supported by grants from the National Cancer (P01-CA108671 ARM) and Associazione Italiana Ricerca Cancro (AIRC IG23525, ARM). Profs. Antonia D’Errico, Ilaria Bassi and Simone Carotti are gratefully acknowledged for providing access to the tissue repositories of St. Orsola Hospital, Bologna and Campus Bio-medico, Rome, Italy.

## Declaration of Interest

The authors declare that the research was conducted in the absence of any commercial or financial relationships that could be construed as a potential conflict of interest. ARM received funding for research from Dompé Pharmaceutics, Forbius, Novartis for other projects.

## Notes

https://zenodo.org/record/6384028#.Yj2NtDfMLzc

https://zenodo.org/record/7974753

### Competing Interest Statement

ARM received funding for research from Dompe Pharmaceutics, Forbius, Novartis.
All the other authors do not have competing interest.

### Summary of Updates

The results and discussion sections were revised for clarity. New information has been added to the following figures: Figure 1 and Figure 2. All the figures and supplementary were improved for clarity.

