## Supplemental figures and tables for "GATA1-defective immune-megakaryocytes as possible drivers of idiopathic pulmonary fibrosis"

**Supplemental item legends**

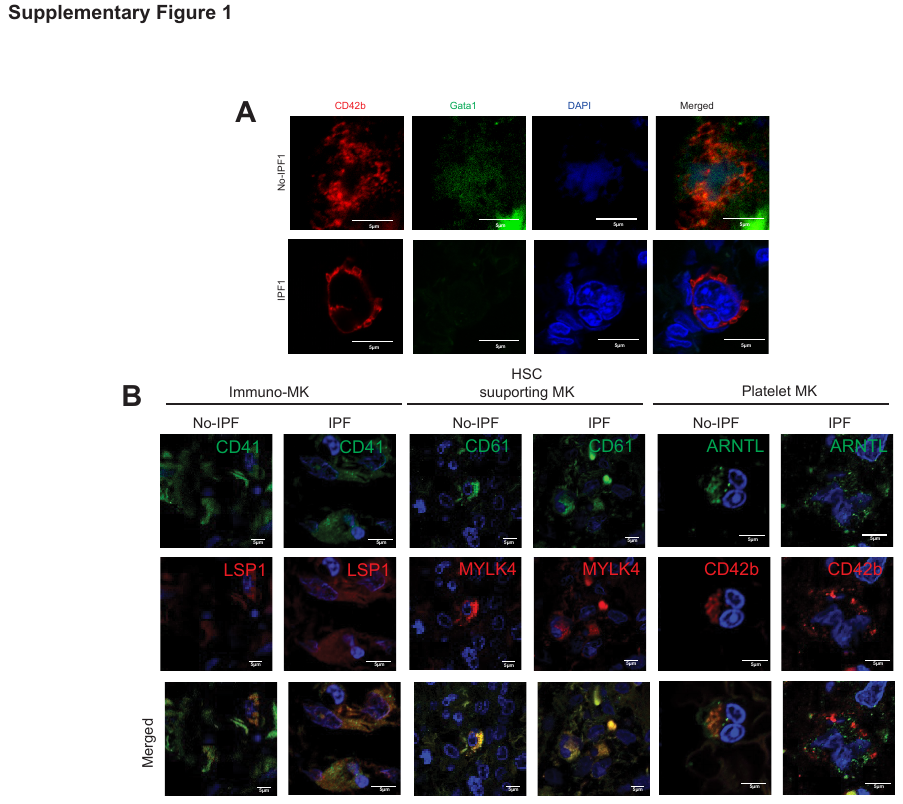

**Figure S1. Signals in the individual channels of confocal analyses of representative human MK.** A) Representative MK in the red (CD42b), green (anti-GATA1) and blue (DAPI) channel showing the detection of GATA1 in the nucleus of the CD42pos cells from the No-IPF lung but not in that from IPF patients. Merged images are presented on the right. Magnification 1200x (see **Figure 1A)** B). Representative images of immune-MK in the green (CD41 green) and red (LSP1) channels; of HSC-supporting MK in the green (CD61) and red (MYLK) channels and of Platelet-poised MK in the red (CD42b) and green (ARNTL) channel from the lung of one No-IPF and one IPF patient, as indicated. DAPI is shown in all the images. The merged signals are presented on the bottom, for clarity. The CD41pos cell from the No-IPF patient is LSP1 neg. Magnification 1200x (see **Figure** **2A**)

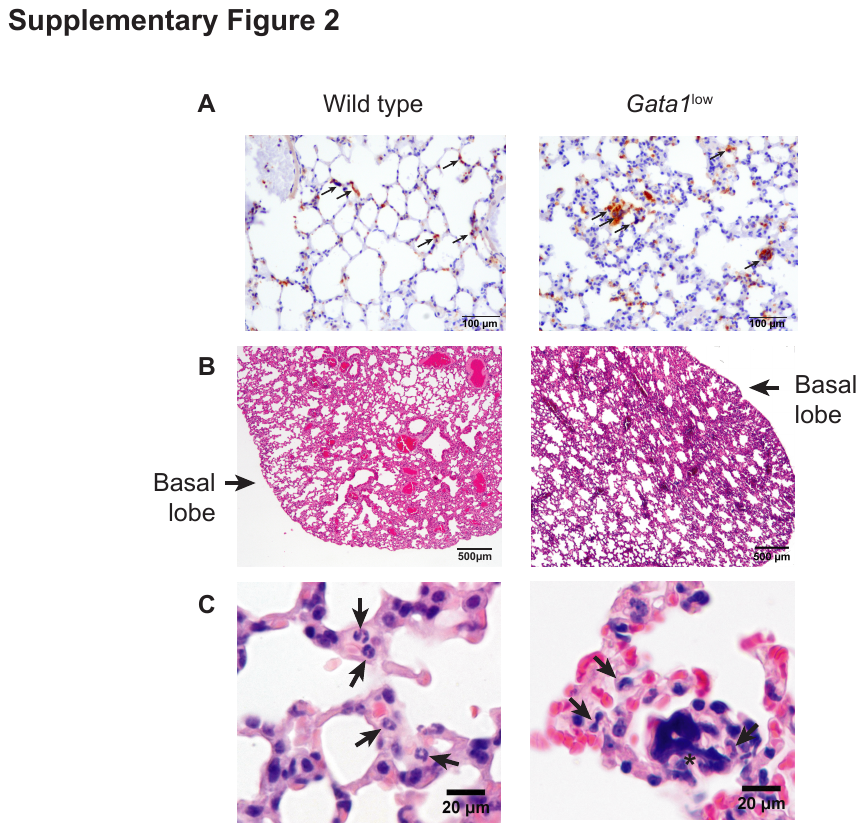

**Figure S2. Localization of the MK and of fibrosis within the architecture of the lung from representative WT and *Gata1^low^* littermates.** A) Images of the lungs from one WT and one *Gata1^low^* mouse indicating that in *Gata1^low^* mice MK are localized within the parenchyma. Magnification 200x. B) Hematoxylin & Eosin staining of the basal lobe of the lung from one WT and one *Gata1^low^* littermate showing that interstitial pneumonia extends from the alveolar septae to the subpleural interstitium in the lung from both WT and *Gata1^low^* littermates (one representative 12 months old mouse out of a total of 12 WT and 22 *Gata1^low^* mice investigated). C) Images at larger magnification of B) showing that lung inflammation is characterized by the presence of neutrophils (the arrows) in the alveolar septae of both WT and *Gata1^low^* mice (the arrows) but that in *Gata1^low^* lungs neutrophils are often localized near a megakaryocyte (the asterisk). Magnification 200x in A); 40x in B) and 400x in C), respectively.

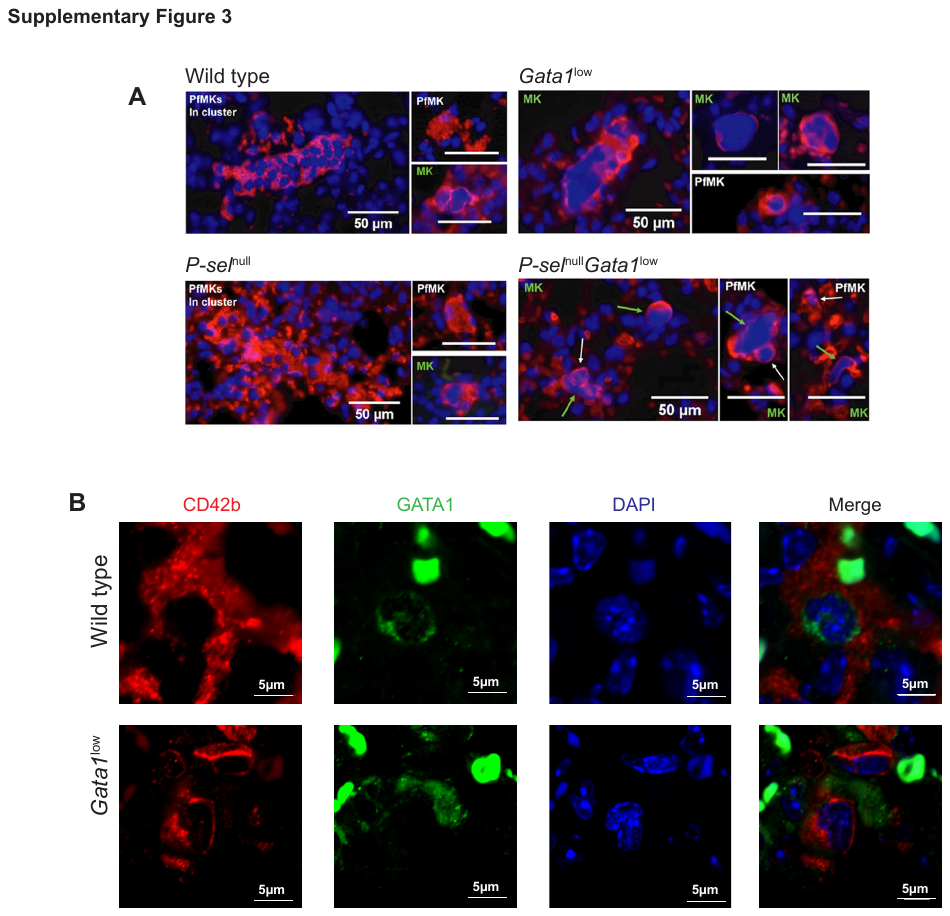

**Figure S3. The megakaryocytes in the lung from old *Gata1^low^* mice have an immature morphology and are hypomorhic for GATA1.** A) Immunofluorescence analyses with the CD42b antibody of lung sections from wild-type (WT) and *Gata1^low^* littermates, as well as from *P-sel^null^* and *P-sel^null^Gata1^low^* mice (all 15-months old), as indicated. Green arrows indicate MK without platelet territories, white arrows indicate mature MK ready to release platelets (PfMK). Magnification 400X. B) Signals in the individual channels (CD42b, red, anti-GATA1, green, and DAPI, blue) of representative MK from WT and *Gata1^low^* mice. Merged images are presented on the right for clarity. Magnification 1200X.

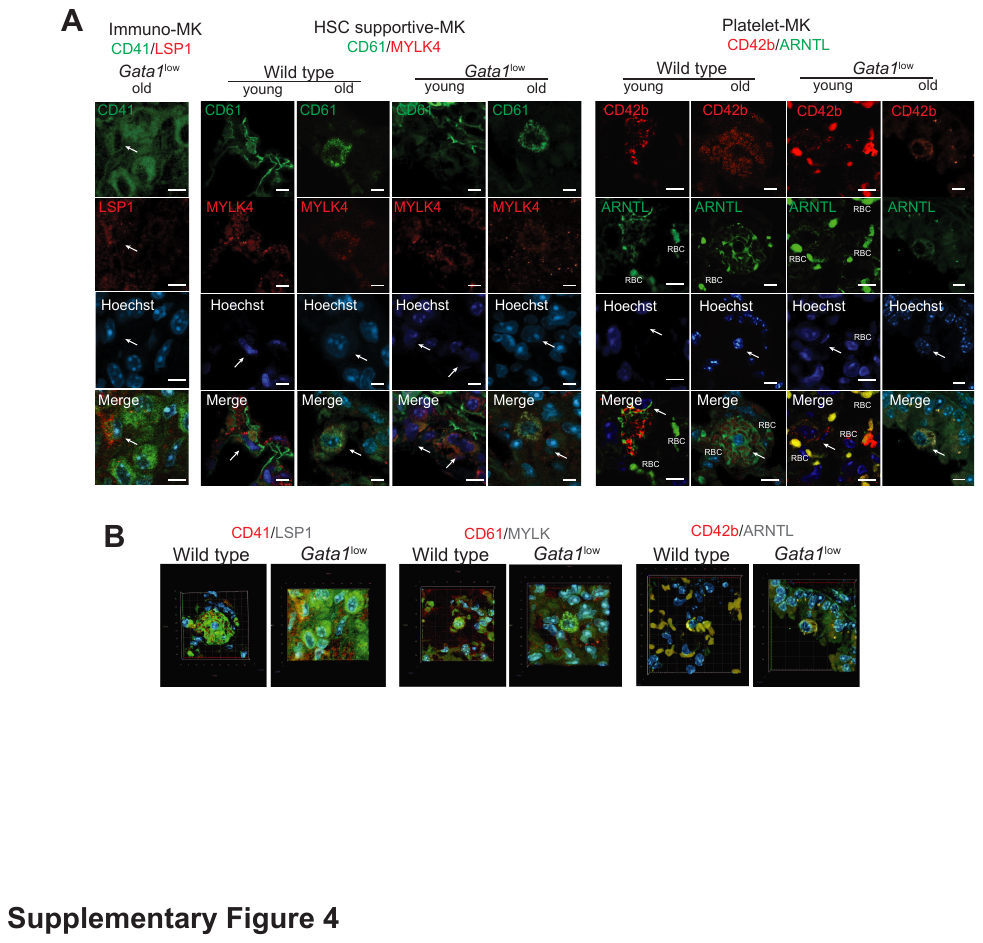

**Figure S4. Megakaryocytes with the phenotype of immuno-MK are detected mainly in the lung from old *Gata1^low^* mice.** A) Signals in the individual channels with anti-CD41 (green) and LSP1 (red)(immune-MK), CD61 (green) and MYLK4 (red)(HSC-supporting MK) and CD42b (red) and ARNTL(green) (Platelet-MK) from the lung of young and old WT and *Gata1^low^* mice, as indicated. Nuclei are stained with DAPI (blue). Merged images are presented on the bottom per clarity. CD41^pos^/LS1^pos^ were not detected in young and old WT and young *Gata1^low^* mice. Magnification 1200x, scale bars 5μm. B) Validation by tridimensional reconstructions of the colocalization of the CD41/LSP1, CD41/MYLK and CD42b/ARNTL signals in megakaryocytes from wild-type and *Gata1^low^* mice as indicated. Magnification 1000x. The corresponding movies may be retrieved at Doi: 10.5281/zenodo.7974752; https://zenodo.org/record/7974753.

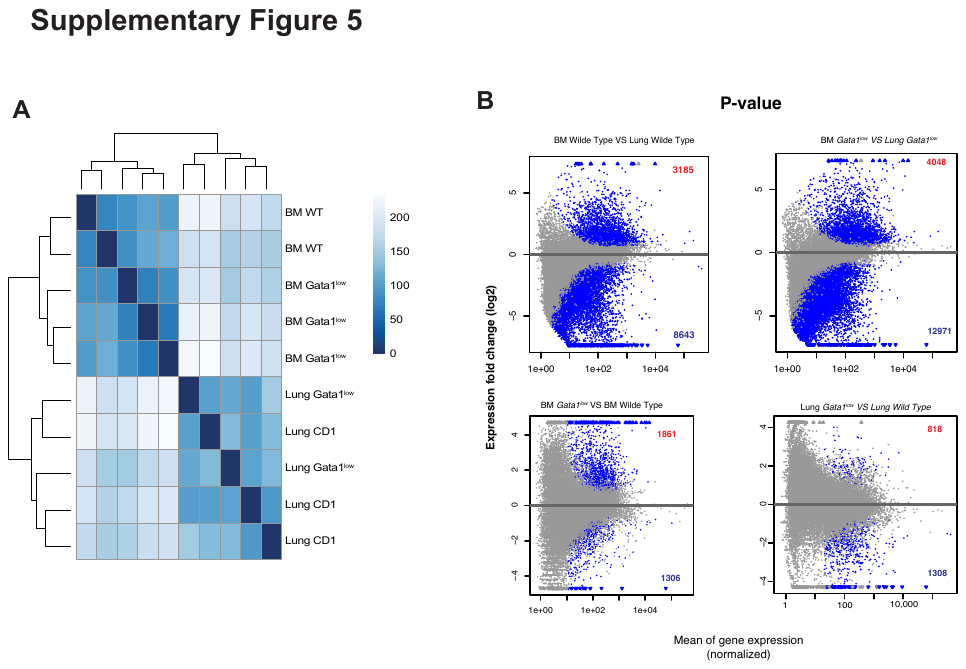
**Figure S5. The megakaryocytes from the lung and bone marrow of old WT and *Gata1^low^* mice have different expression profiling.** A) Heatmap showing the clusterization of the samples, highlighting the difference between the MKs from the lung and from the bone marrow (BM) on both mouse strains. B) MA plots indicating the number of genes expressed at significantly different levels between the megakaryocytes from the bone marrow and those from the lung of WT and *Gata1^low^* mice. The number of genes significant up- and downregulated between groups are indicated in red and blue, respectively.

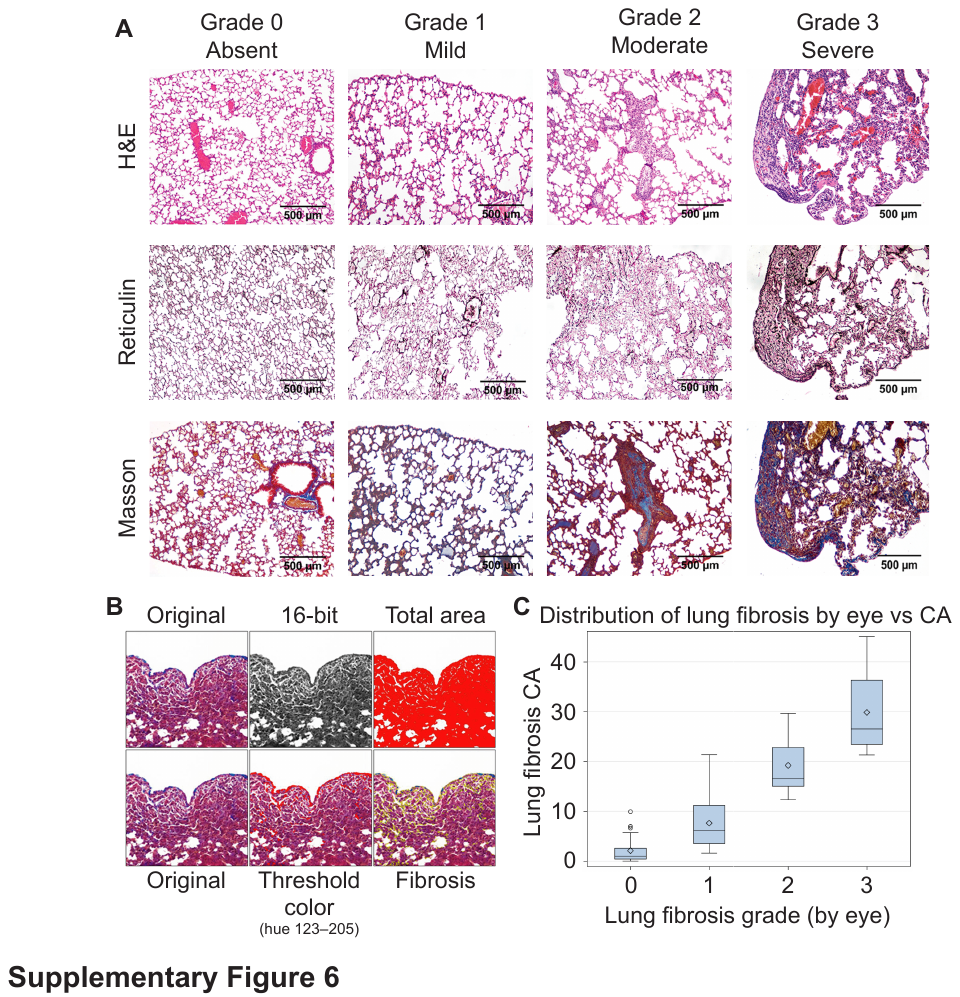

**Figure S6. Semiquantitative and quantitative assessment of fibrosis in the lungs**. Fibrosis was assessed by semiquantitative criteria based on morphological evaluation of the stains by an expert veterinary pathologist (A) and by computer-based quantifications of the intensity of the reticulin or Masson staining (B). For the semiquantitative evaluations in A, the levels of fibrosis were divided into four grades of progressive intensity: Grade 0, presence of collagen fibers only at the bronchovascular level and absence of septal alveolar fibrosis; grade 1, presence of focal mature collagen fibers in the alveolar-septal interstitium; grade 2, deposition of multifocal to coalescing collagen fibers in the alveolar-septa interstitium; grade 3, deposition of collagen bundles that irregularly thicken the alveolar septa and alterations of the overall pulmonary architecture. The panels present hematoxylin & Eosin, Reticulin and Masson's trichrome staining of consecutive sections from the lungs of one representative mouse for each grade of fibrosis. B) Images depicting the process of computer assisted quantification of fibrosis in one lung section stained by Masson’s trichrome. The process includes the determination of the portion of the areas containing collagen fibers (blue channel, Hue 123-205) with respect to the total areas analyzed. C) boxplots showing the distribution of quantitative (0-100% fibrosis) assessment in the lungs from 47 *Gata1^low^* mice divided by semiquantitative (grades 0-3) fibrosis grade. The trend of increasing percent of fibrosis with increasing grade suggests that there is an overall agreement between results obtained with the two methods.

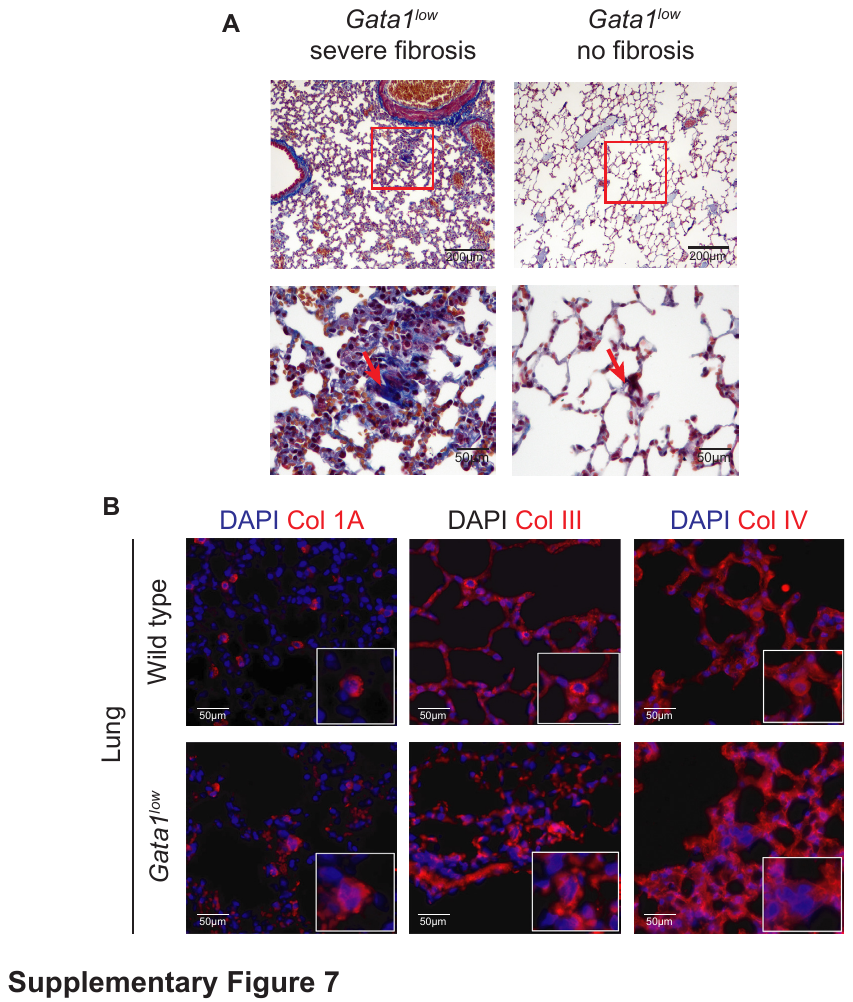

**Figure S7. The megakaryocytes from the lungs of old *Gata1^low^* mice are surrounded by fibers of collagen 1A and III.** A) Masson’s trichrome staining of sections from representative lungs of *Gata1^low^* mice expressing either severe or no fibrosis, as indicated. The images at higher magnification in the rectangles indicate that MK are embedded within the collagen fibers while in that without fibrosis they are located within the alveolae septa. Magnification 100x and 400x in the top and lower panels. B) Immunofluorescence analyses with anti-Col 1A, anti-Col III or anti-Col IV antibodies (red) of lungs from representative WT and *Gata1^low^* littermates. Nuclei were stained with DAPI (blue). The rectangles show representative MK, recognized by morphology, at greater magnification that, in the case of the *Gata1^low^* lung, are surrounded by fibers of Col I and Col III. Collagen IV, used as control, is strongly expressed in both groups, consistent with its known expression along the basal membrane of the alveolar septae.

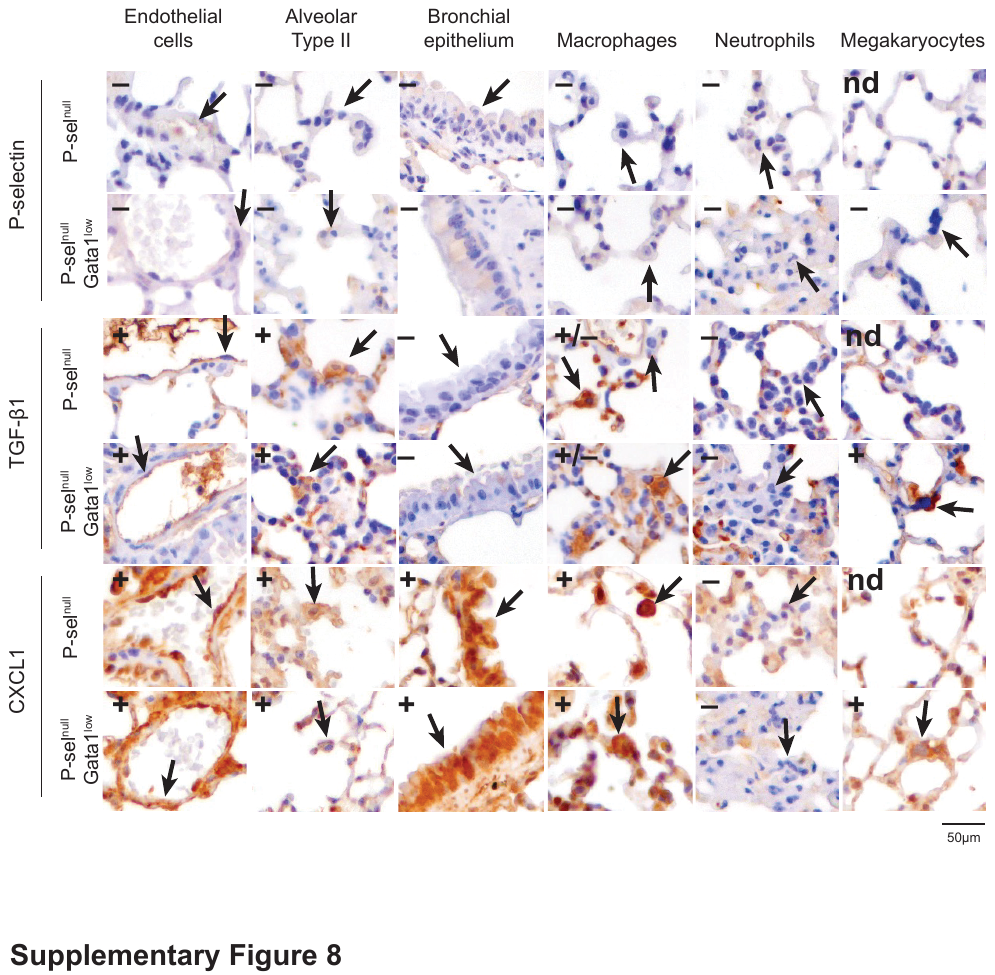

**Figure S8. Genetic deletion of *P-sel* has modest effects on the TGF-β and CXCL1 content of endothelial cells, alveolar type II cells, bronchial epithelium, macrophages and megakaryocytes carrying the *Gata1^low^* mutation (compare these results with those presented in Figure 9).** P-SEL, TGF-β1 and CXCL1 content in endothelial cells, alveolar type II cells, bronchial epithelium, macrophages, neutrophils and megakaryocytes (indicated by the arrow) from representative *P-sel^null^* and *P-Sel^null^/Gata1^low^* littermates (both 13-months old), as indicated. Cells were identified according to the morphological criteria described in the legend of **Figure 9**. Magnification 400x. Protein content is quantitatively assessed as undetected (-) and detected (+). nd= cell type not detected.

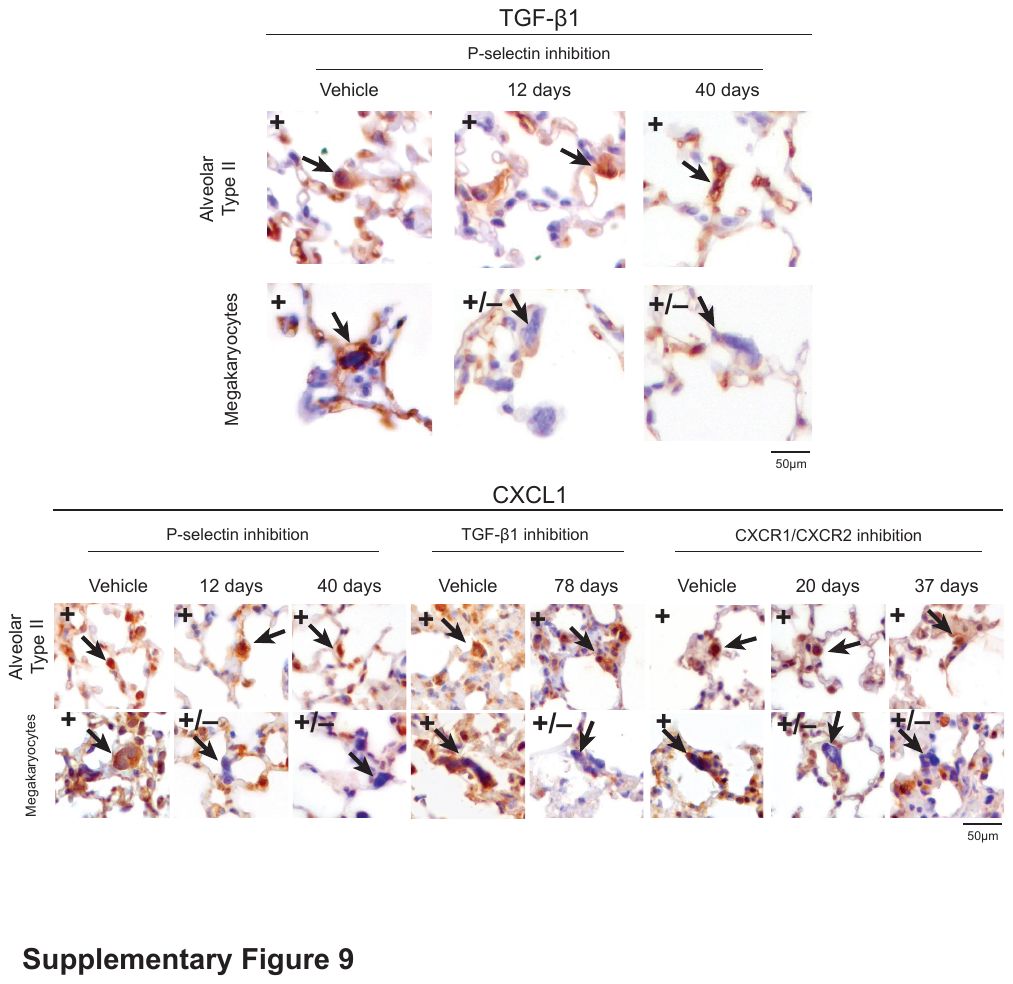

**Figure S9. Treatment with the P-SEL inhibitor reduced the CXCL1 content of both megakaryocytes and alveolar type II cells and TGF-β content in megakaryocytes while that with the TGF-β or CXCR1/2 inhibitor reduced only the CXCL1 content in megakaryocytes.** TGF-β1 (A) and CXCL1 (B) content in alveolar Type II cells and in MK (indicated by the arrow) from representative *Gata1^low^* mice treated either with vehicle or the P-SEL, the TGF-β1 or the CXCL1 inhibitor, as indicated. Cells were identified according to the morphological criteria described in the legend of **Figure 9**. Magnification 400x. Protein content is quantitatively assessed ad undetected (-), barely detectable (+/-) and detected (+).

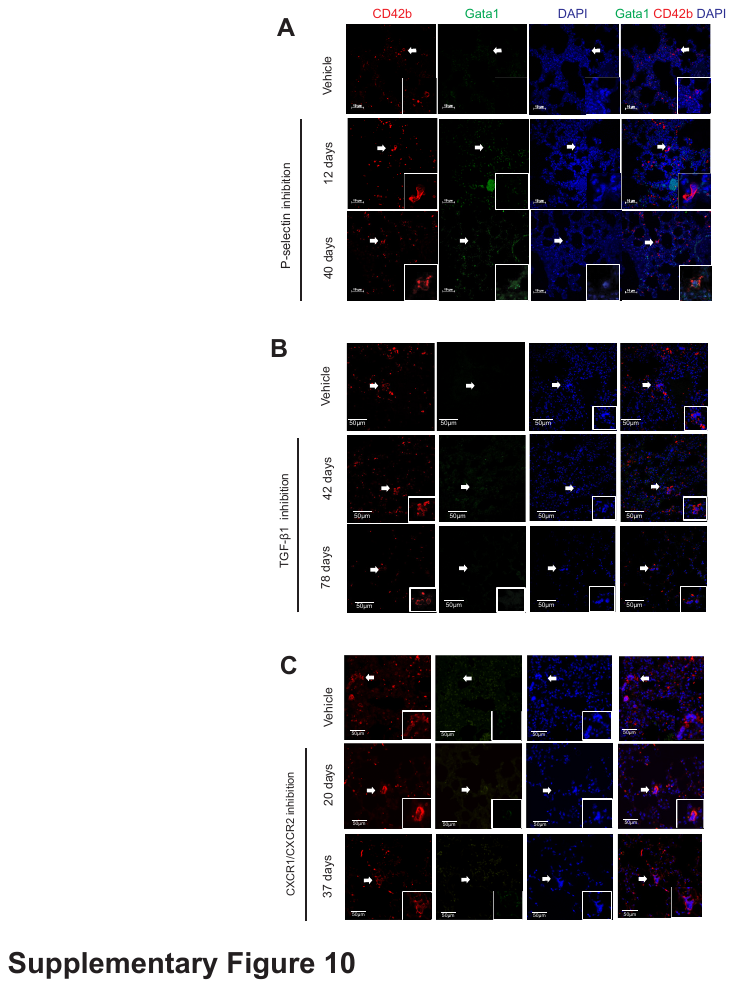

**Figure S10. Signals in the individual channels (CD42b, red, anti-GATA1, green, and**

**DAPI, blue) of the merged images of the MK presented in Figure 13A, C and E.** The merged signals are also presented for clarity. Magnification 400x

**Supplementary Tables**

**Table S1**. Clinical detail of the human subjects included in the study.

| **No** | **Age** |  | **Diagnosis** | **Origin** |
| --- | --- | --- | --- | --- |
| 11 | 76 | Control No-IPF1 | Lung Cancer | Tissue Bank, Campus Biomedico Hospital, Rome, Italy |
| 12 | 73 | Control No-IPF2 | Lung Cancer |  |
| 13 | 76 | Control No-IPF3 | Lung Cancer |  |
| 14 | >60 | IPF1 | IPF | Tissue Bank, Ospedale S. Orsola, Bologna, Italy |
| 15 | >60 | IPF2 | IPF |  |
| 16 | >60 | IPF3 | IPF |  |

| **Table S2.** Comparison of histological parameters between WT and *P-sel*^null^ | | | | | |
| --- | --- | --- | --- | --- | --- |
|  | Genotype | |  |  |  |
|  | *P-sel*^null^ (N=16) | WT (N=18) | Unadjusted  p-value | Age-adjusted  p-value | Sex-adjusted  p-value |
| **Age (months)** |  |  |  |  |  |
| N (Missing) | 16 (0) | 18 (0) |  |  |  |
| Mean (±SD) | 9.4 (±4.60) | 8.7 (±5.56) | 0.67^1^ |  |  |
| Median | 10.0 | 10.5 |  |  |  |
| Range | 3.0, 16.0 | 2.0, 18.0 |  |  |  |
| **Sex**, n (%) |  |  |  |  |  |
| F | 5 (41.7%) | 5 (31.3%) | 0.70**^b^** |  |  |
| M | 7 (58.3%) | 11 (68.8%) |  |  |  |
| N (Missing) | 12 (4) | 16 (2) |  |  |  |
| **Total MK/144mm^2^** |  |  |  |  |  |
| N (Missing) | 4 (12) | 9 (9) |  |  |  |
| Mean (±SD) | 4.8 (±3.60) | 3.6 (±2.42) | 0.49**^a^** | 0.51**^c^** |  |
| Median | 3.7 | 3.0 |  |  |  |
| Range | 1.9, 10.1 | 0.0, 3.8 |  |  |  |
| **Small MK/144mm^2^** |  |  |  |  |  |
| N (Missing) | 4 (12) | 9 (9) |  |  |  |
| Mean (±SD) | 4.4 (±3.92) | 3.4 (±2.53) | 0.57**^a^** | 0.60**^c^** |  |
| Median | 3.4 | 3.0 |  |  |  |
| Range | 0.9, 10.0 | 1.0, 8.3 |  |  |  |
| **Large MK/144mm^2^** |  |  |  |  |  |
| N (Missing) | 4 (12) | 9 (9) |  |  |  |
| Mean (±SD) | 4.9 (±4.82) | 0.3 (±0.42) | 0.01**^a^** | 0.001**^c^** |  |
| Median | 5.0 | 0.0 |  |  |  |
| Range | 0.5, 9.1 | 0.0, 1.2 |  |  |  |
| **Severity of lung inflammation**, n (%) |  |  | >0.99**^b^** | 0.97**^d^** | 0.93**^d^** |
| 0 | 3 (23.1%) | 3 (25.0%) |  |  |  |
| 1 | 5 (38.5%) | 4 (33.3%) |  |  |  |
| 2 or 3 | 5 (38.5%) | 5 (41.7%) |  |  |  |
| N (Missing) | 13 (3) | 12 (6) |  |  |  |
| **Semiquantitative fibrosis**, n (%) |  |  | >0.99**^b^** | 0.73**^d^** | 0.15**^d^** |
| 0 | 8 (61.5%) | 8 (66.7%) |  |  |  |
| 1 | 4 (30.8%) | 4 (33.3%) |  |  |  |
| 2 or 3 | 1 (7.7%) | 0 (0.0%) |  |  |  |
| N (Missing) | 13 (3) | 12 (6) |  |  |  |
| **Quantitative fibrosis** |  |  |  |  |  |
| N (Missing) | 13 (3) | 12 (6) |  |  |  |
| Mean (±SD) | 4.2 (±4.60) | 3.0 (±3.10) | 0.47**^a^** | 0.49**^c^** | 0.08**^c^** |
| Median | 2.1 | 1.9 |  |  |  |
| Range | 0.4, 16.4 | 0.0, 9.0 |  |  |  |
| **^a^**Equal variance two-sample t-test p-value; **^b^**Fisher Exact p-value; **^c^**Parameter p-value from multiple linear regression with age or sex as covariate; or **^d^**Parameter p-value from ordinal logistic regression with age or sex as covariate | | | | | |
